# PALS1 is a key regulator of the lateral distribution of tight junction proteins in renal epithelial cells

**DOI:** 10.1101/2023.04.27.538411

**Authors:** Ann-Christin Groh, Simon Kleimann, Pavel Nedvetsky, Malina Behrens, Annika Möller-Kerutt, Verena Höffken, Sujasha Ghosh, Uwe Hansen, Michael P. Krahn, Alexander Ludwig, Klaus Ebnet, Hermann Pavenstädt, Thomas Weide

**Author notes:** corresponding author Thomas Weide, PhD University Clinics of Muenster (UKM), Internal Medicine D (MedD), Albert-Schweitzer-Campus 1, Building A14, 48149 Muenster, Germany.

## Abstract

The evolutionarily conserved Crumbs (CRB) polarity complex, which consists of the core components CRB3a, PALS1 and PATJ, plays a key role in epithelial cell-cell contact formation and cell polarization. Recently we observed that deletion of one *Pals1* allele in mice results in functional haploinsufficiency characterized by renal cysts. To address the role of PALS1 at the cellular level, we generated PALS1 knockout MDCKII cell lines using the CRISPR/Cas9 system. The loss of PALS1 resulted in increased paracellular permeability indicative of an epithelial barrier defect. This barrier defect was associated with a redistribution of several tight junction-associated proteins from bicellular cell-cell contacts to tricellular junctions. The regulation of tight junction protein localization at bicellular junctions by PALS1 was dependent on its interaction with PATJ. Together, our data uncover a critical role of PALS1 in the correct positioning of tight junction proteins to bicellular junctions.

## INTRODUCTION

Apicobasal cell polarization establishes an asymmetric distribution of lipids and proteins in the plasma membrane and forms the basis for the physiological role of numerous epithelia in mammalian organs. Studies of the last two decades in *Drosophila melanogaster*, in the zebrafish *Danio rerio* and in mammalian cell cultures demonstrated the essential role of the evolutionarily conserved Crumbs (CRB) complex in epithelial cell polarity development and epithelial tissue morphogenesis [1, 2]. The CRB core complex consists of the name-giving type I transmembrane protein Crumbs (in mammals mainly CRB3a), LIN7 (lin-7 homolog C), PALS1 (protein associated with LIN7), and PATJ (PALS1-associated tight junction protein) [2, 3]. Recently, it has been shown that the CRB complex components CRB3a, PATJ and PALS1 localize just apical of the tight junction in a novel compartment called the vertebrate marginal zone (VMZ) [4, 5]. *In vitro* studies identified an essential role of PALS1 in the formation of tight junction (TJ) and adherens junctions (AJ), providing evidence that cell junction assembly and cell polarization are closely connected biological processes [5–7].

Cell polarization and the formation of cell contacts are of outstanding importance for mammalian kidneys, particularly for renal nephron epithelia. The physiological function of various tubular nephron epithelia depends on the asymmetric distribution of numerous channels and transporters to regulate the resorption and recycling of nutrients, as well as on the formation of TJ to regulate the salt and ion homeostasis and to concentrate the primary ultra-filtrate into excretable urine.

Several studies indicate an important role for the CRB complex in kidney function and imply that dysfunction of CRB core components might acts as a causative or aggravating factor for renal diseases. For instance, mice lacking CRB3 or LIN7 develop cysts in the renal tubules of the kidneys [8–10]. In addition, heterozygous PALS1 *knockout* mice with a reduced PALS1 expression in all nephron epithelia [11] developed proteinuria and formed numerous cysts leading to death within the first two months after birth [11, 12]. Remarkably, in case of PALS1 the loss of only one allele was sufficient to cause this phenotype, indicating a crucial and dose-dependent function of PALS1 in renal epithelial cells [11]. However, it remains unclear if reduced PALS1 levels lead to defects in cell polarization, or rather affect the formation of cell contacts between the epithelial cells of the nephron.

In this study, we analyzed the role of PALS1 in the formation of AJs and TJs in Madin Darby canine kidney (MDCKII) cells in which both *PALS1* alleles were inactivated using CRISPR/Cas9. These cells show moderate defects in cell polarization but exhibit dramatically increased paracellular permeability. Strikingly, several TJ components such as ZO-1 and Occludin as well as the PALS1 binding partner PATJ are lost from bicellular TJs in *PALS1* KO cells. Thus, our data provide novel evidence that PALS1 regulates the barrier function in MDCKII cells by controlling the lateral distribution of TJ components.

## Methods

### Constructs and Cloning

We generated pENTR constructs with human PALS1 cDNA inserts (Gene ID: 64398) encoding for PALS1 full length (aa 1-675) and two PALS1 deletion mutants (ΔECR: aa 118-674, ΔL27N: aa 178-675) using the pENTR^TM^ Directional TOPO® Cloning Kit (Invitrogen). The pENTR constructs were shuttled into modified pQCXIH-GW-GFP or pQCXIH- GW-Halo plasmids using the L/R reaction of the Gateway^TM^ Cloning system (Invitrogen), according to the manufacturer’s instructions. All primers and oligonucleotides are listed in suppl. Table ST1.

### Generation and establishment of PALS1 knockout cell lines

To generate a PALS1 knockout in MDCKII cells (#ECACC 00062107) frameshift mutations were introduced via CRISPR/Cas9. CRISPOR was used to design the following guide RNA/DNA-sequences CACCGATCATTAGTCGGATAGTAAA, AAACTTTACTATCCGACTAATGATC. The gRNAs (100µM, 8µl each) were phosphorylated and annealed via adding 2µl 10x PNK buffer, 1 µl ATP (5mM) and 1 µl Polynucleotide Kinase and incubation for 1 h at 37°C followed by incubation at 95°C for 5 min to stop the reaction. Resulting oligonucleotides were inserted into the px459 plasmid via digestion using BbsI and ligation by T4 Ligase. MDCKII cells with a confluency of 60% were transfected using Lipofectamine™ 2000 following the manufacturer’s protocol. Selection was done using puromycin (4 µg/ml) for 48 h. Potential knockout cells were separated to analyze single cell clones. Potential clones were sequenced and only clones with frameshift mutations were kept. Heterozygous clones were further analyzed after cloning with Zero Blunt® TOPO® PCR cloning kit to separate individual alleles.

### Cultivation of MDCKII cell lines

MDCKII cell lines were grown in modified Eagle’s medium (MEM, Lonza) containing 5% fetal calf serum (FCS) and 1% 100x Penicillin/Streptomycin/L-Glutamine (P/S/G). HEK293T cells were grown in DMEM medium containing 10% FCS and 1% P/S/G. The cells were cultivated at 37°C and 5% CO2. For transient transfection, cells were transfected using Lipofectamine 2000 according to the manufacturer’s manual.

MDCKII cells were seeded on coverslips, on ThinCert™ cell culture inserts (0.4 mm-pore transwell filters, Greiner Bio-one) to reach full polarization, or in a matrix hydrogel (basement membrane extract, BME, from R&D Systems) in 3D to form cysts. Medium was exchanged every other day. On coverslips cells were grown up to a confluency of 100%, in transwell filters for 7-10 days and in BME for 5-7 days before using them for immunofluorescence staining. In case of mixed monolayers, wildtype and KO cells were mixed 1:1 and seeded on coverslips or ThinCert™ cell culture inserts, respectively.

### Generation of stable cell lines

Stable cell lines were generated via retroviral gene transfer, as described earlier[13]. In brief, HEK293T cells were transiently transfected with the helper plasmids pCMV-VSV-G and pUMVC and pQCXIH plasmids encoding for full length PALS1 or deletion mutants fused to HALO- or EGFP fluorescence tags. Supernatant containing the virus particles were added to MDCKII WT and KO cells after being filtered through a sterile 0.45 µm filter (Millipore). Here, one volume of DMEM including supplements was mixed with one volume of virus-containing medium supplemented with polybrene (8µg/ml). After 24h, virus-containing medium was collected and stored at 4°C. Cells were left in medium without virus to regenerate for 24h. After regeneration period, the second round of transduction using the same virus-containing medium was performed for another 24h. After another 24 for regeneration, transduced cells were selected using hygromycin (250 µg/ml). All established cell lines were tested for a constitutive overexpression of PALS1 proteins by live cell imaging, or Western blot.

### Generation of PALS1 shRNA lines

Short hairpin RNAs (shRNAs) were purchased as complementary oligonucleotides that when annealed contained the *HindIII* (5’) and *BamHI* (3’) sticky ends for cloning into the pSuper vector. The PALS1 shRNA B oligonucleotides had the following sequence: 5’-CC<u>GGAGATGAGGTTCTGGAAAT</u>TCAAGAG<u>ATTTCCAGAACCTCATCTCC</u>TTTTT-3’. All plasmids were verified by Sanger sequencing. The shRNA-transfected MDCKII cells were grown in 200 µg/ml hygromycin to select for stable clonal cell lines. The PALS1 rescue line was generated by transfecting shRNA-resistant PALS1-GFP into the clonal PALS1 shRNA line B1, followed by clonal selection in 400 µg/ml geneticin [5]. PALS1 shRNA and PALS1-GFP rescue IF and WB experiments were performed as described [5].

### TEER measurements

MDCKII cells were seeded on ThinCert™ cell culture inserts (12-mm, 0.4 mm-pore transwell filters, Greiner Bio-one) to measure the transepithelial electrical resistance (TEER). The electrical resistance between the apical and the basolateral chamber was measured using a two-electrode resistance system. To calculate the unit area resistance (Ω x cm^2^), the electric resistance was multiplied by the growth area of the transwell filters (1.13 cm^2^). The measurement was performed every 5 min from the moment of seeding the cells to 7 days after seeding. Medium was changed once after 3-4 days. Values were normalized to the wildtype control (WT) values.

### Generation of 3D cysts and permeability assay

To grow 3D MDCKII cysts, ibidi µ-Slide 8-well chambers were coated with 8µl BME. For solidification of the BME coated wells were incubated at 37°C with 5% CO2 for 15-30 min. Cells were detached from a 2D monolayer using incubation at 37°C with 5% CO2 in 10x Trypsin/EDTA solution until reaching a single cell suspension. Cells were seeded (5,000 cells/cm^2^), resuspended in 250 µl in medium containing 10% FCS, 1 % P/S/G and 2.5 % BME. Cells were cultured for 5-7 days at 37 °C with an atmospheric CO2 concentration of 5%. Medium was exchanged every second day. Cysts were either used for immunofluorescence staining (7 days) or for 3D permeability assays (5 days).

To measure the permeability of cysts, fluorescently labeled dextran of different sizes was used (Alexa647, 10k; FITC, 3-5k). Medium was exchanged with 200 µl live cell imaging medium (15mM HEPES, ad HBSS). Cysts were imaged live using the HC PL APO 40x/1.10 W motCORR CS2 objective of a confocal microscope (TCS SP8, Leica Biosystems). Cysts were imaged every minute for 15-20 min in total. After imaging time point t0, medium was supplemented with 50 µl dextran solution (10 µM Alexa647, 12.5 µM FITC). Cyst permeability was quantified by signal intensity comparing the lumen intensity against the average intensity outside of the cyst.

### Transmission electron microscopy (TEM)

MDCK cysts cultivated in cell culture plates were washed with PBS and then fixed in 2% (v/v) formaldehyde and 2.5% (v/v) glutaraldehyde in 100 mM cacodylate buffer, pH 7.4, at 4°C overnight. After washing in PBS, specimens were postfixed in 0.5% (v/v) osmium tetroxide and 1% (w/v) potassium hexacyanoferrate (III) in 0.1 M cacodylate buffer for 2 h at 4°C followed by washing with distilled water. After dehydration in an ascending ethanol series from 30% to 100% ethanol, specimens were two times incubated in propylene oxide each for 15 min. The cells were detached from the surface of the cell culture plate during the propylene oxide treatment. Finally, detached cells were embedded in Epon using beem capsules. Ultrathin sections were cut on an ultramicrotome and collected on copper grids. Electron micrographs were taken at a Phillips EM-410 electron microscope using imaging plates (Ditabis, Pforzheim, Germany).

### Preparation of cell lysates and Western blot analyses

Cells were grown on 6-well plates up to a confluency of 95%, washed twice with 1x PBS and lysed with 300 µl 1x SDS sample buffer [20% (v/v) Glycerin, 125 mM Tris-HCl pH 6.8, 10% (w/v) SDS, 0.2% (w/v) Bromphenol blue, 5% Mercaptoethanol, ad 1L H2O]. For homogenization, samples were passed through a blunt 20 G needle (0.9 mm diameter) and heated up to 95°C. Lysates were centrifuged for 1 min at 14000 x g and stored at -20°C until usage. Western blot analysis was performed as previously described ^2^. First, cell lysates were boiled at 95 °C for 5 min. Proteins were separated via SDS-PAGE using 10% gels (BioRad). Then, proteins were transferred to a PVDF membrane (Millipore) following by incubation in blocking powder (5 % milk powder in TBST [0.05 % Tween 20]) for 1h at room temperature (RT). Probes were incubated with primary antibodies (suppl. Table ST2) diluted in BSA (2.5 g BSA powder in 50 ml TBST) o/n at 4 °C. Afterwards, the membrane was washed three times with TBST followed by incubation with horseradish peroxidase coupled secondary antibodies (suppl. Table ST3) for 1h at RT. The membrane was washed again for three times with TBST before a chemiluminescence detection reagent (Clarity; BioRad) was added and the membrane was imaged using an Azure Biosystems imager (c600; Azure Biosystems).

### Immunofluorescence analysis

Cells on coverslips and transwell filter insets were grown either as pure cultures or as co-cultures of two different cell lines mixed in a ratio of 1:1. 3D cysts were grown as described before. All cells were washed twice with cold 1x PBS and fixed in 4% Paraformaldehyde (PFA) for 20 min at RT. After cells were washed three times with 1x PBS, they were quenched for 10 min in 50 mM NH4Cl followed by another three rounds of washing in 1x PBS. Then, cells were permeabilised in 0.2 % Triton X-100 for 5 min. After washing the cells three times in IF wash buffer [0.2 % (w/v) gelatin, 10 % (v/v) 10x PBS, 0.2 % (v/v) Triton X-100, ad H20)] and blocked [90 % (v/v) IF wash buffer, 10 % (v/v) normal goat serum (NGS) for 20 min at RT. Cells were incubated o/n in antibody dilutions buffer (98 % (v/v) IF wash buffer 2 % (v/v) NGS, containing specific antibodies (suppl. Table ST2). Next, cells were washed three times with IF wash buffer and transferred into fluorophore-conjugated secondary antibodies (suppl. Table ST3) solved in antibody solution containing DAPI (1:5,000). After an incubation of 30 min at RT, cells were washed the last time with 1x PBS and were carefully rinsed in dH2O before either mounted in 6-12 µl mowiol on object slides (coverslips, ThinCert™ cell culture inserts) or in IMM mounting medium (ibidi GmbH, Cysts). Samples were imaged on a confocal microscope (TCS SP8, Leica Biosystems) using a 40x water objective, or a spinning disk microscope (CorrSight, Thermofisher) using a 40x or 63x oil objective (NA 1.3 or 1.4; Zeiss Plan Apochromat). Confocal stacks were processed, and analyses were performed with FIJI.

### Quantification and statistical evaluation

Images were analyzed with Fiji (https://fiji.sc/). Movies were created using LAS X (Leica Biosystems). Graphs were created and statistical analysis was performed with the GraphPad PRISM software. Data show the mean ± the standard error of the mean (SEM). The value of n and the definition is indicated in the corresponding figure legends and results. For comparison of two groups the Mann-Whitney U test was used, for comparison of more than two groups the Kruskal-Wallis-test was performed corrected for multiple testing with the Dunn’s correction (*p<0.05, **p<0.01, ***p<0.001). If not indicated differently, all data represent at least three independent repeats.

## Results

The knockout of PALS1 in MDCK cells causes a severe permeability defect. To ensure the complete absence of PALS1 we established MDCKII KO cells by CRISPR/Cas9 and selected a target sequence within the exon 7 of the canine *PALS1* gene, encoding the PALS1 PDZ domain (suppl. Fig. S1A). The two knockout clones KO1 and KO2 showed no expression of the two PALS1 splice variants (Fig. 1A). Immunofluorescence (IF) analyses confirmed the complete absence of PALS1 at cell-cell junctions (Fig. 1B). Since WT1 and WT2 as well as KO1 and KO2 show identical phenotypes, from now on only data from WT1 (WT) and KO1 (KO) are presented as representative of each phenotype.

**Figure 1:**
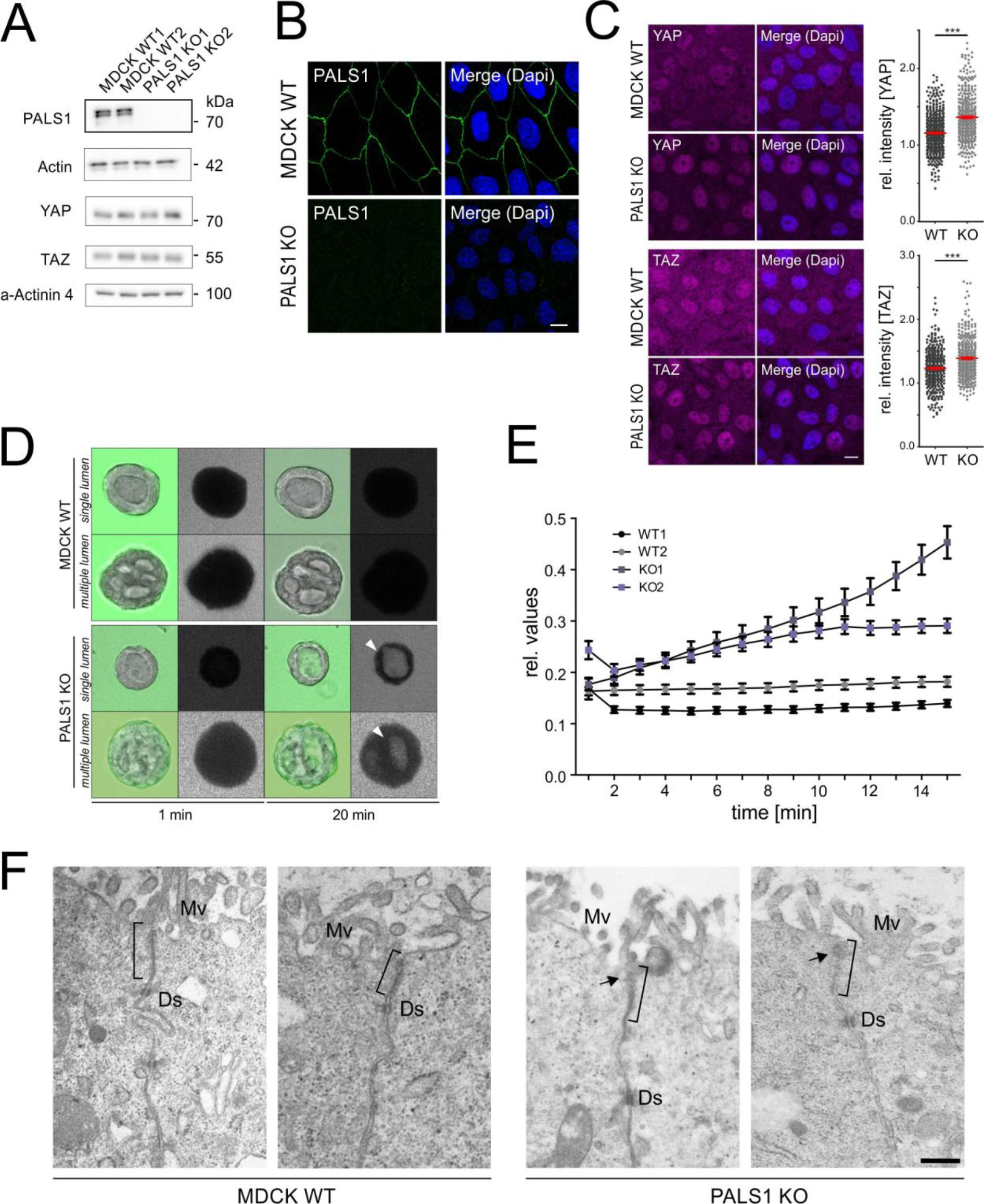
MDCK PALS1 knockout cell lines show a severe permeability defect. (A) Western blot analyses of MDCKII wildtype (WT) and PALS1 knockout cells (KO) clones: Lysates of MDCKII wildtype cells WT1 or cells that passed through the entire CRISPR/Cas9 process without changing the *PALS1* gene sequence (WT2) and both PALS1 KO clones (KO1 and KO2) were analysed for expression of PALS1 and Hippo target proteins YAP and TAZ. Actin and α-Actinin-4 served as loading controls. No PALS1 expression was detectable in PALS1-deficient samples. Scale bar: 10 µm. (B) Immunofuorescence (IF) analyses of MDCK WT and MDCK PALS1 KO cell lines. In contrast to the starting pool of MDCKII cells (WT1) and a clone that showed no mutation after the CRISPR/Cas9 procedure (WT2), PALS1 KO cells show the complete absence of PALS1 at cellular junctions. Scale bar: 10 µm. (C) IF analyses of Hippo target proteins YAP and TAZ. Nuclei of MDCK PALS1 KO cells show a significant nuclear accumulation of YAP and TAZ. For quantification (graphs on the right) the mean signal intensity within each nucleus was measured and compared to the mean signal of the whole cells (cytoplasm and nucleus). The YAP quantification based on the evaluation of n=423 WT and n=370 KO cells of three independent experiments (N=3). TAZ quantification (N=3) was done with n=378 WT and n=359 KO cells. ***≙ p ≤ 0.001. (D) Permeability assays. Labelled dextran (4 kDa) was used to analyze the permeability of PALS1 KO and WT MDCK cysts. No uptake was detectable in either single- or multiple-lumen WT cysts. However, the signal intensity was increased in single and multiple-lumen Pals1 KO cysts. (E) Time course of the permeability assays (see F, N=3) for cysts (n=50) derived from two independent WT (WT1, WT2) and PALS1 KO (KO1, KO2) cell lines. Signal intensity (relative value) analysis over time within the lumen normalized to the surrounding signal in the medium. (F) TEM images of apical junctional regions (black parentheses) of MDCK WT (left) and PALS1 KO (right). In MDCK WT cells, electron dense and clearly structured apical junctional regions which contain the tight junctions were found, whereas in PALS1 KO cells, this region is less electron-dense and more diffuse (arrows). Ds: desmosome. Mv: microvilli, scale bar 1 µm

The CRB complex was identified as an upstream regulator of the Hippo pathway[14]. We recently demonstrated that loss of PALS1 in the kidney affects YAP/TAZ signaling *in vivo*. Therefore, we tested, how the complete loss of PALS1 affects the intracellular localization of the Hippo transcriptional co-activators YAP and TAZ. To address this, we performed quantitative IF studies with MDCK WT and PALS1 KO cells. The IF studies showed an accumulation of YAP as well as TAZ in nuclei of PALS1 KO cells, indicating a direct correlation between the loss of PALS1 from junctions and the nuclear presence of YAP and TAZ (Fig. 1C).

Next, we tested the trans-epithelial-electrical-resistance (TEER) of PALS1 KO cells. Compared to the wild-type cell lines, the TEER values of both KO cell lines were strongly reduced. In addition, a time delay of the maximum TEER values (peak) was observed in the KO cells (suppl. Fig. S1B), similar to previous studies [6].

MDCK cells embedded in extracellular matrix (ECM) form 3D spherical cysts with a single apical lumen. These cysts are surrounded by a layer of apicobasally polarized cells. 3D MDCK cysts share many properties with *in vivo* epithelia and are an excellent *in vitro* model for studying polarity and cell-cell contact formation and integrity[15]. Therefore, we analyzed MDCKII WT and PALS1 KO cells in 3D cyst formation assays. Both WT and PALS1 KO cells formed cysts with single-, multiple and no-lumen. However, multiple- and no-lumen cysts were more prevalent in PALS1 KO cell lines (suppl. Fig. S1C), similar to PALS1 knockdown MDCK cell lines[6]. We stained cysts with antibodies against ZO-1, a marker for TJs, and E-Cadherin as a marker for AJs. In the cells along the cysts, the ZO- 1 signal shows mainly an apical distribution facing the lumen. E-Cadherin, on the other hand, is mainly found basolaterally between the individual cells of the cysts (suppl. Fig. S1D). This is the case in WT as well as in Pals1 KO cells and can be observed in single lumen cysts as well as in cysts with multiple lumens.

The altered TEER dynamics of PALS1 KO cells grown in 2D monolayers (suppl. Fig. S1B) suggest differences in the formation or functionality of cell junctions. Therefore, we next used 3D cultures to investigate if PALS1 influences the barrier properties of cell-cell contacts in highly polarized cells. To address this aspect, accumulation of labeled dextran in cysts lumens were analyzed over time. Cysts of PALS1 KO cells showed an accumulation of labelled dextran in their lumen (Fig. 1D, suppl. Fig. S2A). The effect was visible within the first two minutes, increased over time and was also observed in cysts forming multiple lumens (Fig. 1E, suppl. Fig. S2B).

Next, we analyzed the cell-cell junctions in 3D cysts for potential changes in their structure by transmission electron microscopy (TEM). The apical regions of cell-cell contacts, which contain the TJs appeared more diffuse and less electron dense in PALS1 KO cells compared to WT cells. In contrast, desmosomal structures showed no obvious differences between WT and PALS1 KO cells in 3D cysts, indicating that PALS1 regulates the structure of cell-cell junctions in the apical region of epithelial cells (Fig. 1F).

PALS1 depletion results in altered lateral distribution of tight junction proteins at bicellular junctions. The increased permeability of cell-cell contacts in PALS1 KO cysts and the ultrastructural changes in the apical region of cell contacts of KO cells found by TEM argue for changes in TJ organization and function. To test this, we first investigated the distribution of the TJ components ZO-1, Occludin, Claudin1 and JAM-A, as well as the AJ protein E-Cadherin in WT and PALS1 KO cells in 2D monolayers. MDCKII WT cells showed a lateral “chicken-wire-like” distribution of E-Cadherin and ZO-1 (Fig. 2A, upper panel). In PALS1 KO cells ZO-1 disappeared from bicellular junctions, leading to a “dot-like” pattern of ZO-1 that was caused by strong ZO-1 signals at the tricellular junctions (Fig. 2A, lower pattern). The pattern of E-Cadherin remained unchanged in PALS1 KO cells. A similar pattern was found for the TJ protein Occludin. Like ZO-1, it showed a reduced intensity along bicellular junctions in PALS1 KO cells (suppl. Fig. S3A). By contrast, JAM-A, a protein that is also enriched in TJ [16, 17], showed a normal “chicken-wire-like” distribution in MDCK WT and PALS1 KO cells (suppl. Fig. S3B).

**Figure 2:**
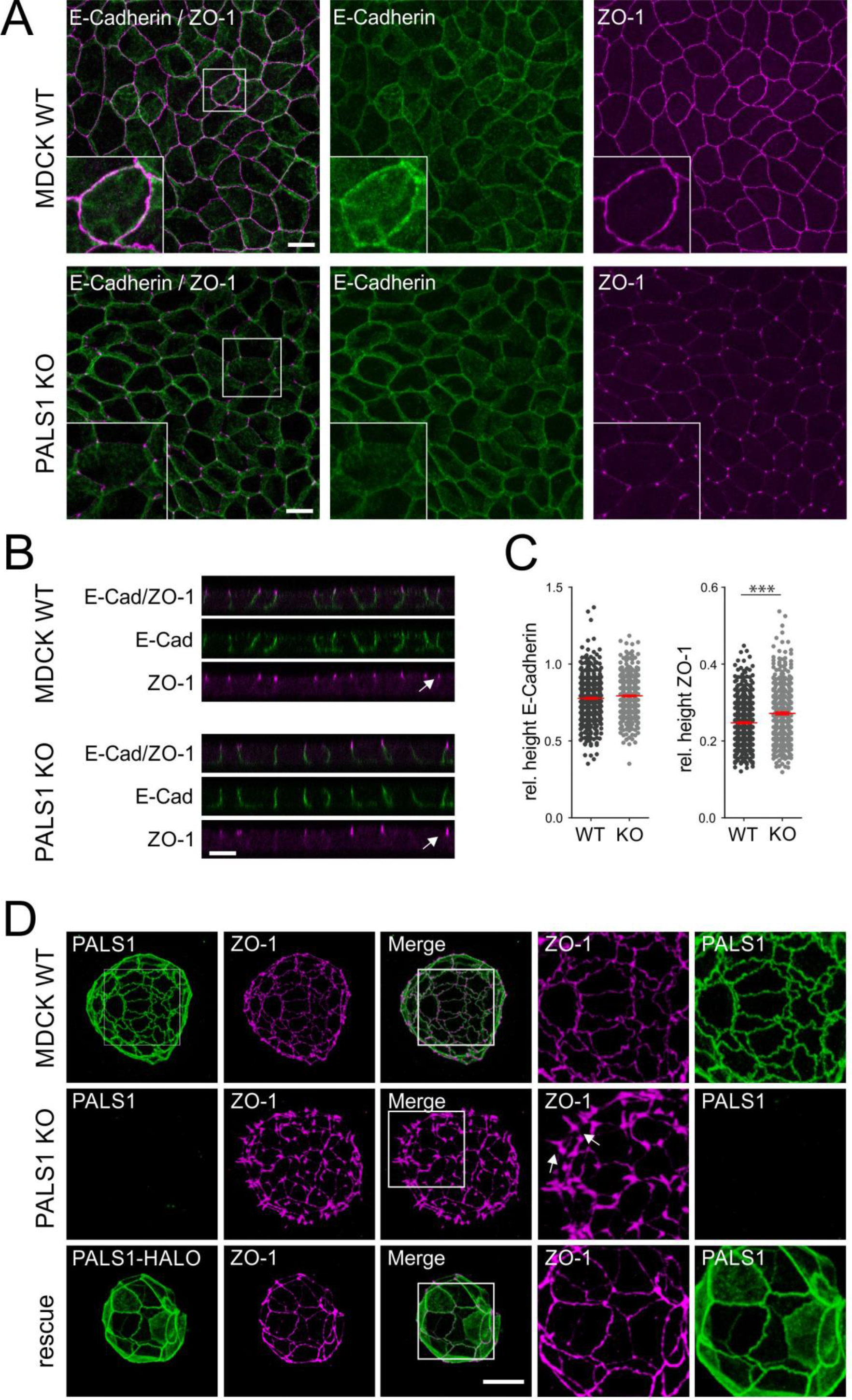
PALS1 depletion alters the lateral distribution of the TJ protein ZO-1 (A) IF analyses using E-Cadherin and ZO-1. The distribution of E-Cadherin (green) showed no obvious differences in 2D cultures of WT and PALS1 KO MDCK cells. In contrast, the distribution of ZO-1(magenta) in PALS1 KO cells shows a strong decrease of the signal in bicellular contacts, which is accompanied by an accumulation of this TJ protein in tricellular contacts. Scale bar: 10 µm. (B) ZO-1 and E-Cadherin were analyzed in vertical direction (z-axis) in polarized WT and PALS1 KO cells. For the analysis, the distribution length of the ZO-1 (WT: n=544; KO: n=400) and E-Cadherin (WT: n=527; KO: n=487) signal was measured and normalized to the full height of the cell monolayer. Scale bar: 10 µm. (C) The relative length of the E-Cadherin signal shows no significant change between WT and PALS1 KO cells (left graph). The relative length of ZO-1 the signal area is significantly increased in PALS1 KO cells in comparison to WT cells (right graph). ***≙ p ≤ 0.001. Mean with SEM indicated. (D) IF analyses of 3D MDCK cysts. In comparison to WT cysts, cysts from PALS1 KO show a “spike-like” distribution of ZO-1 (magenta, arrows), accompanied by a discontinued distribution of the ZO-1 signal at bicellular junctions. PALS1 KO cysts that express full length PALS1 fused to a HALO tag (PALS1-HALO) lose their “spike-like” ZO-1 misdistribution phenotype. Brightness and contrast were increased equally for all images. Scale bar: 10 µm

After observing the strong decrease of TJ proteins at bicellular junctions, the distribution of ZO-1 and E-Cadherin were analyzed in vertical direction (z-axis) in polarized WT and PALS1 KO cells (Fig. 2B). For the analysis, the length of the ZO-1 and E-Cadherin signals were measured and normalized to the full height of the cell monolayer. The analysis revealed no overt changes in the distribution of E-Cadherin in polarized WT and PALS1 KO monolayers. However, the ZO-1 distribution was altered in monolayers of PALS1 KO cells as a significant elongation of the signal along the z-axis was observed (Fig. 2C).

We next compared 3D cultures of WT and PALS1 KO cells. In cysts, PALS1 deficiency resulted in ZO-1 positive “spike-like” elongations at tricellular junctions accompanied by a discontinuous ZO-1 pattern in bicellular junctions. These spikes were absent in the MDCKII WT control cysts (Fig. 2D). To verify the specificity of the CRISPR/Cas9-mediated PALS1 KO we re-expressed HALO-tagged PALS1 in PALS1 KO cell lines. PALS1 KO cysts that expressed PALS1-HALO fusion proteins re-established the continuous ZO-1 distribution along bi- and tricellular junctions in 3D. “Spike-like” elongations at tricellular junctions were not observed (Fig. 2D). Thus, a proper PALS1 expression is required to regulates the continuous distribution of ZO-1 in 2D and 3D cultures of epithelial cells.

The continuous lateral ZO-1 distribution is directly linked to cellular PALS1 expression. To investigate if PALS1 has an impact on the distribution of TJ proteins in adjoining cells, we used 1:1 mixed monolayers of MDCKII WT and PALS1 KO cells and performed IF studies with antibodies against PALS1, ZO-1 (Fig. 3A/B), E-Cadherin, Occludin and Claudin1 (suppl. Fig. S3). At bicellular contacts, the highest signal intensity of ZO-1 was observed between junctions of WT cells (WT-WT). Cell junctions between one WT and one KO (WT-KO) cell showed reduced signal intensities and the contacts of two KO cells (KO-KO) showed the lowest levels. In comparison to the expected stepwise decrease of the PALS1 signal (WT-WT versus WT-KO and KO-KO, respectively) the reduction in the ZO-1 signal in WT-KO and KO-KO contacts was moderate, with average values not below 80% of the WT-WT contacts (Fig. 3C). IF-studies of co-cultures grown on glass coverslips confirmed our observations on transwell filters and again showed a reduction in TJ proteins (ZO-1, Occludin, and Claudin 1) at bicellular junctions (suppl. Fig. S4). In contrast, tricellular junctions shared by KO cells (KO-KO-KO) showed the strongest ZO-1 signal intensity. The intensity in mixed junctions (WT-WT-KO and WT-KO-KO, respectively) was between the values of the homogenous tricellular junctions composed only of WT or KO cells (Fig. 3D). Together the data suggest that the continuous distribution of ZO-1 in bicellular pools is directly linked to the PALS1 expression level within a cell contact.

**Figure 3:**
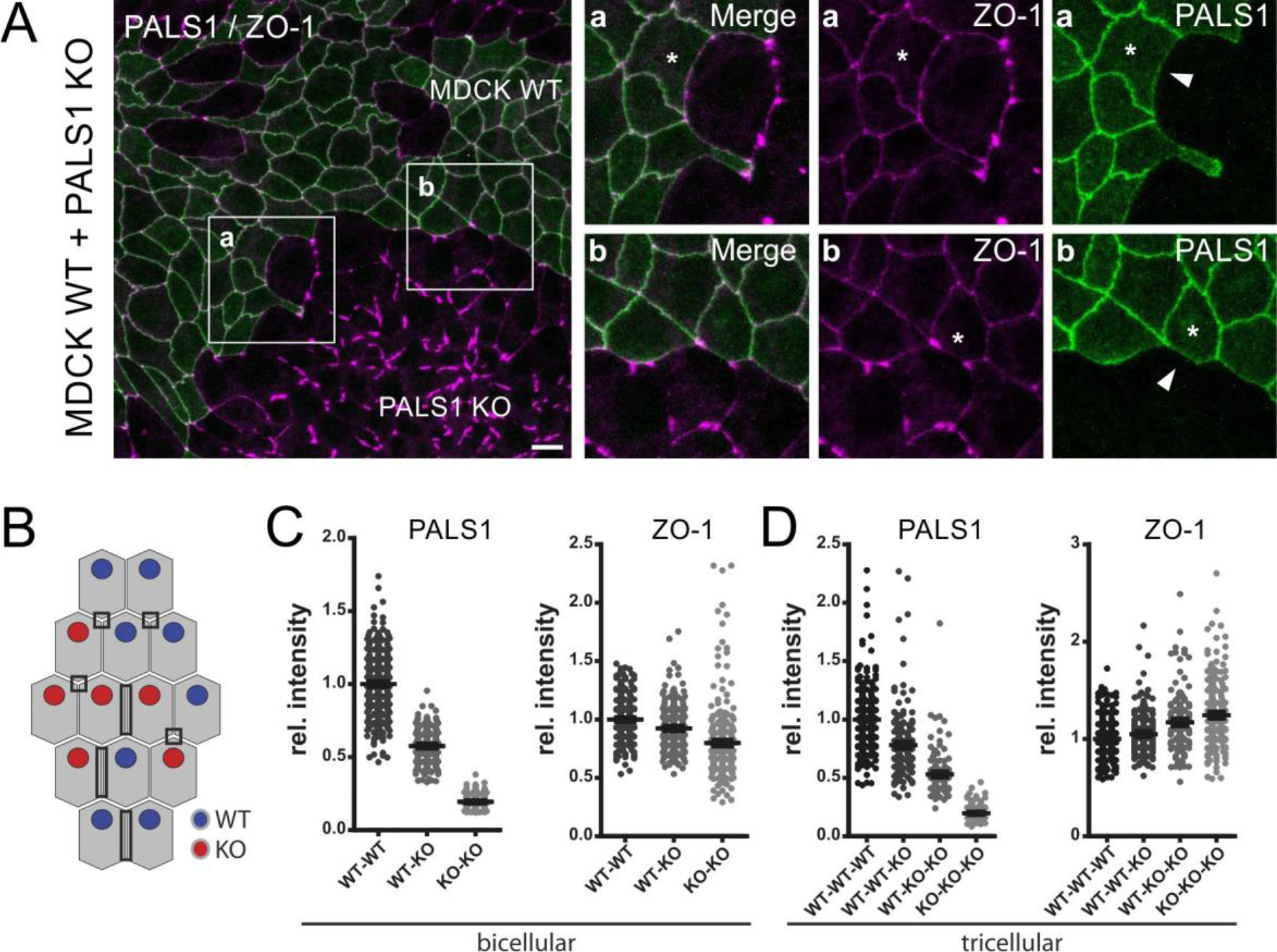
The continuous lateral ZO-1 distribution is directly linked to cellular PALS1 expression. (A) Distribution of PALS1 (green) and ZO-1 (magenta) in 1:1 mixed co-cultures of WT and PALS1 KO MDCK cells. Images show a maximum intensity projection. Brightness and contrast were increased equally for all images. Scale bar: 10µm (B) Scheme of randomly mixed WT (blue) and PALS1 KO cells (red). Mixed monolayers establish different types of bi- and tricellular contacts. Bicellular contacts include three possible variants (WT-WT, WT-KO, and KO- KO) and tricellular contacts four (WT-WT-WT, WT-WT-KO, WT-KO-KO, and KO-KO-KO). (C/D) Quantitative analyses of the relative intensity at bi- and tricellular junctions for either PALS1 or ZO-1. Signals were measured using a region of interest (ROI) with a defined size. Intensities were normalized to the mean intensity of bi- and tricellular junctions of the wildtype (WT-WT and WT-WT-WT), respectively. (C) Quantitative analyses of bicellular junctions. The relative intensity of PALS1 and ZO-1 revealed a significant decrease in signal intensity from WT-WT (n=374) over WT-KO (n=221) to KO-KO (n=270) contacts. All possible cell-cell contact formation were tested for significance and show p-values below p ≤ 0.001 (***). (D) Quantitative analyses of tricellular junctions. The relative intensity of PALS1 is significantly decreased. The more KO cells are part of a tricellular junction the lower the PALS1 signal. *Vice versa*, ZO-1 signal significantly increases in case that at least PALS1 KO cell contributes to a tricellular junction and is strongest in tricellular junction that completely lack PALS1 (D, right graph). All possible cell-cell contact formation were tested for significance and show p-values below p ≤ 0.001 (***), except for WT-WT-WT (n=414) *versus* WT-WT-KO (n=214) and WT-KO-KO (n=136) *versus* KO-KO-KO (n=214) combinations.

The lateral distribution of ZO-1 depends on the presence of PALS1 at tight junctions. Next, we wondered whether membrane targeting of PALS1 controls the distribution of ZO-1. To address this, we expressed two PALS1 mutants with a C-terminal GFP-tag in PALS1 KO cells (Fig. 4A): One mutant lacked the N-terminal evolutionarily conserved region (ECR), which mediates binding to PAR6 (ΔECR: aa 118-675) [18–20]. The other mutant additionally lacked the N-terminal L27 domain (L27N), which binds to PATJ (ΔECR+L27N: aa 178-675) [21]. Full length PALS1 fused to GFP (PALS1-GFP WT) was used as a positive control. Transient transfection of constructs encoding these PALS1- GFP fusion proteins in PALS1 KO cells showed that only GFP-tagged PALS1 WT and the ΔECR deletion mutant can bind to cell junctions and were able to re-establish a continuous ZO-1 distribution. The ΔECR+L27N deletion mutant, lacking the first 177 aa, failed to bind to TJs and showed a diffuse cytoplasmic localization (Fig. 4B).

**Figure 4:**
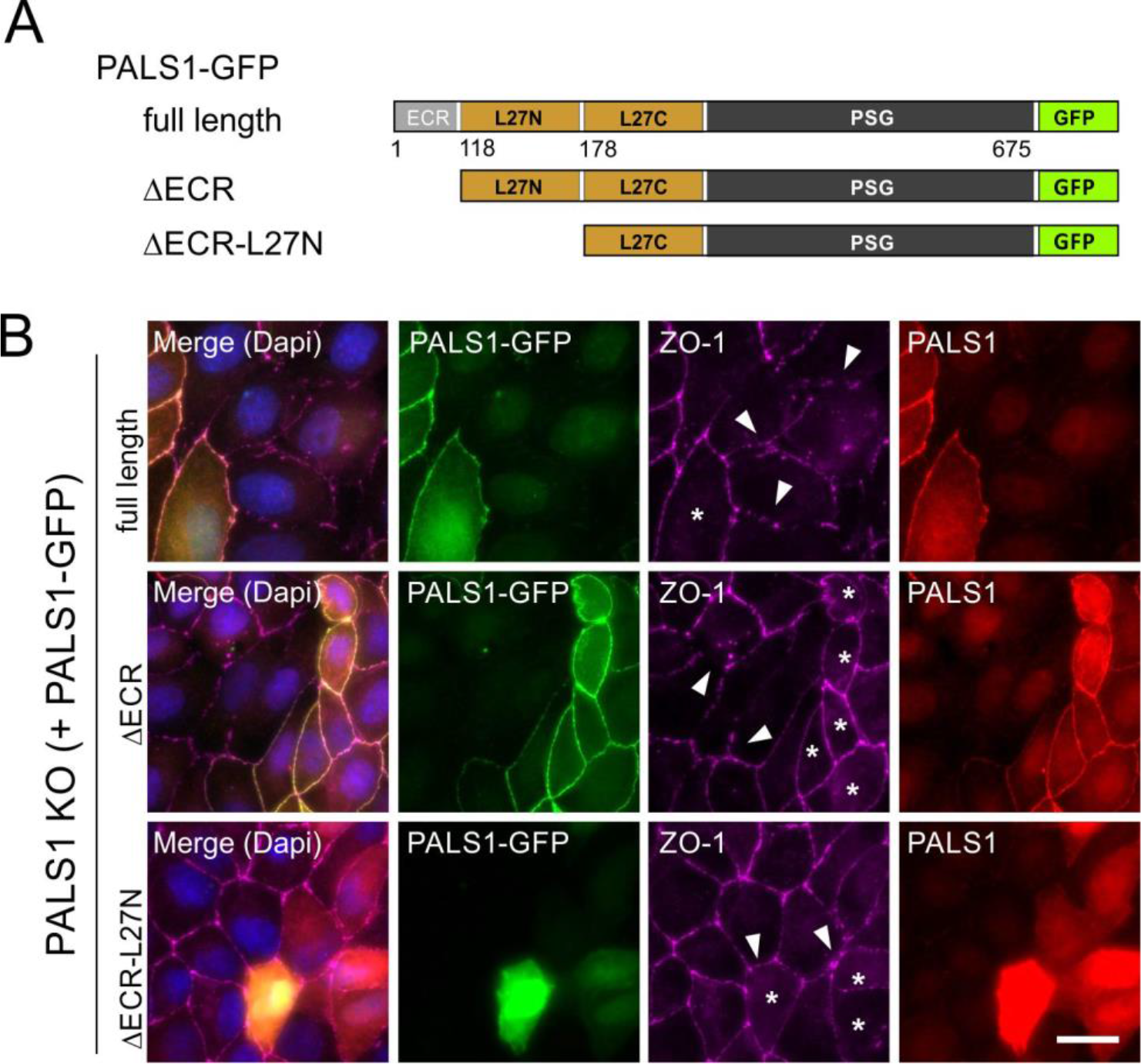
The lateral distribution of ZO-1 requires the PALS1 L27N domain. (A) Schematic of different GFP-tagged PALS1 deletion mutants. MDCK WT cells (B) and PALS1 KO cells (C) were transiently transfected with constructs encoding either the GFP-tagged full length PALS1 (first panel), the ΔECR deletion mutant (lacking first 117 amino acids), and the deletion lacking the first 177 amino acids including theL27N domain (ΔECR+L27N). (B) IF analyses of transiently transfected PALS1 KO cells. GFP-tagged PALS1 full length and ΔECR deletion mutants (green, asterisks) showed binding to membranes co-localize with endogenous ZO-1 (magenta) and can rescue the disrupted discontinuous ZO-1 distribution (magenta). Cells that express the ΔECR+L27N mutant (green, asterisks) maintained the “dot-like” ZO-1 pattern (lowest panel, arrowheads). Scale bar: 20 µm

Both, PALS1 membrane recruitment [21] as well as the PALS1-dependent control of the lateral continuous ZO-1 distribution required the L27N PATJ binding module. This raised the question to what extent PALS1 also determines the intracellular localization of PATJ. We addressed this by IF studies and found that junctional PATJ levels were strongly reduced in PALS1 KO cells, indicating that PALS1 regulates the localization of PATJ. In addition, the remaining PATJ pools in PALS1 KO cells showed a “dot-like” pattern, that strongly overlapped with that of ZO- 1 (Fig. 5A). Next, we analyzed PALS1 KO cells expressing the PALS1-HALO fusion protein. Patches with expression of the PALS1-HALO fusion protein re-established the continuous distribution of endogenous PATJ in bi- and tricellular junctions as well as the lateral distribution of ZO-1 (Fig. 5B). Bicellular contacts between PALS1-HALO positive cells and cells without a detectable PALS1-HALO expression maintained a disrupted ZO-1 distribution, similar to what we had observed in mixed cultures (Fig. 3). This confirmed that PALS1 only on one side of cell-cell contacts is unable to trigger full TJ formation in adjoining cells (Fig. 5C).

**Figure 5:**
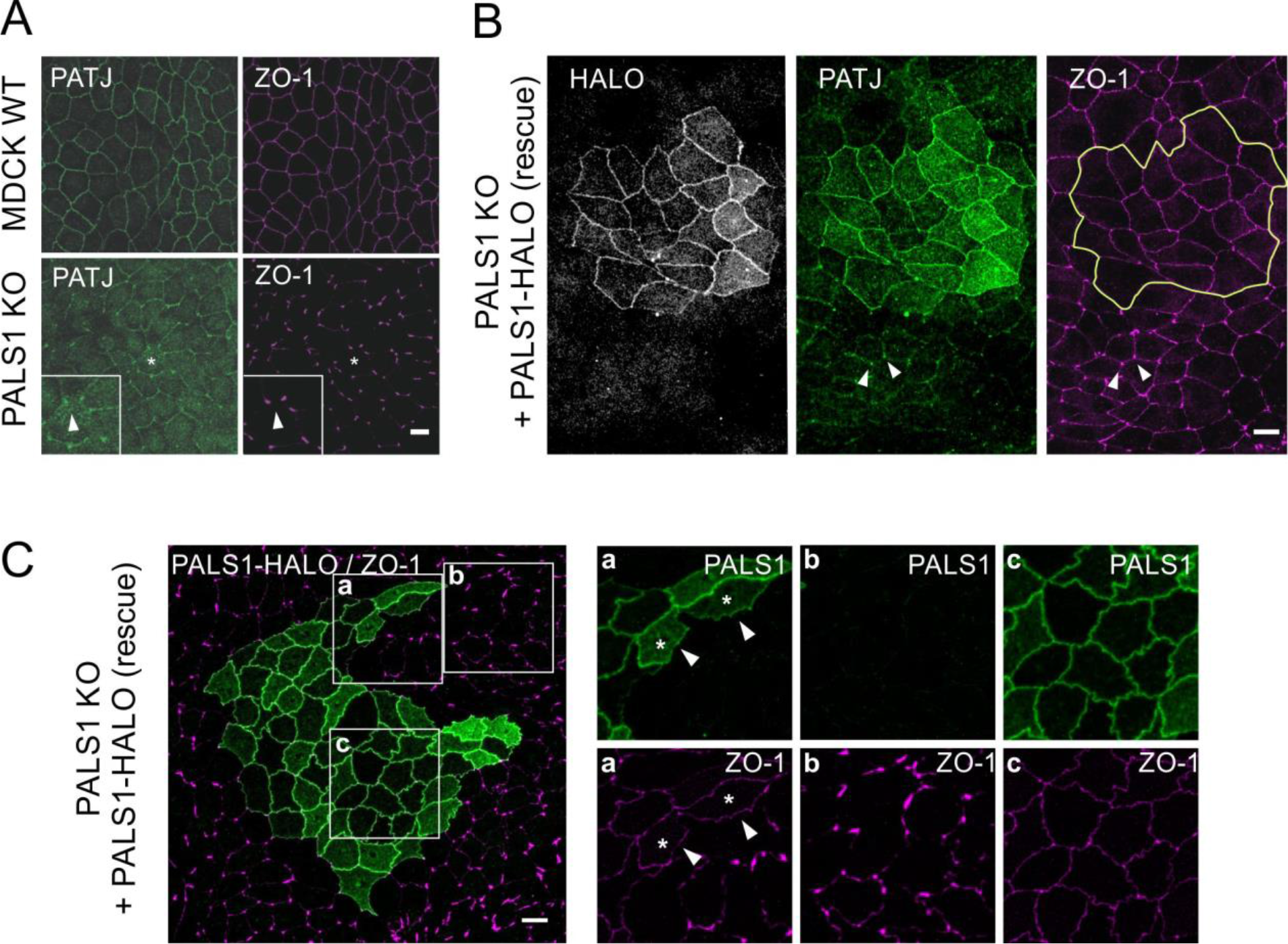
PALS1 controls the lateral distribution of PATJ at cell junctions. (A) IF of WT and PALS1-KO MDCKII cells. *Upper panel*: PATJ (green) and ZO-1 (magenta) strongly co-localize and show a “chicken-wire-like” distribution in MDCK WT cells. In PALS1 KO cells (lower panel) PATJ and ZO-1 levels are reduced in bicellular junctions accompanied by an increased signal intensity in tricellular junction, resulting in a “dot-like” distribution pattern. (B). IF studies of PALS1 KO cells expressing a PALS1-HALO fusion protein. Patches expressing the Pals1-HALO fusion proteins are targeted to TJs in PALS1 KO cells (left). Expression of the PALS1- HALO fusion proteins re-established the expression and TJ associated localization of endogenous PATJ (middle panel) as well and the continuous distribution of ZO-1 (magenta). The yellow line mark cells with PALS1-HALO expression. (C) IF analyses of polarized PALS1 KO MDCKII cells (lower panel) plus cell rescued by the expression of a PALS1-HALO. Cells expressing the PALS1-HALO rescue construct (green patches) show a complete restoration of the ZO-1 “dot-like” mislocalization phenotype of PALS1 KO cells. Detail a: border area between rescued and non- rescued PALS1 monolayer; detail b: non-rescued area; detail c: full-rescued area. Scale bars: 10 µm

The previous experiments (Figures 2-5) demonstrate the central role of PALS1 in controlling the lateral distribution of PATJ and TJ proteins. To clarify to what extent these effects are dose-dependent, we analyzed PALS1 KD cell lines stably expressing a PALS1 shRNA. PALS1 KD cells showed a strong reduction of the PALS1 level (Fig. 6A/B, suppl. Fig. S5). Remarkably, residual PALS1 in PALS1 KD cells showed a weak punctate pattern different to the continuous belt-like staining seen in WT MDCKII cells (Fig. 6A, increased gain). In PALS1 KD cells TJ proteins such as ZO-1, and in particular ZO-3, were strongly reduced along bicellular junctions (Fig. 6C/D). This led to a “dot-like” pattern that was different from the “chicken-wire-like” distribution of TJ proteins ZO-1 and ZO-3 in control cells expressing a non-targeting shRNA (Fig. 6C/D, suppl. Fig. S5D). Co-cultures of PALS1 KD cells and MDCK cells expressing a control shRNA showed a strong decrease in PATJ and Occludin levels at bicellular TJs (Fig. 6E/F). PALS1 KD cells stably expressing a shRNA-resistant PALS1-GFP fusion protein (suppl. Fig. S6A) resulted in an increase of the endogenous PATJ expression (Fig. 6E-G) and in a re-distribution of PATJ along bicellular junctions (suppl. Fig. S6B). Together, these data identified PALS1 as a crucial factor for the continuous lateral distribution of PATJ and a range of bona-fide TJ proteins at bicellular TJs.

**Figure 6:**
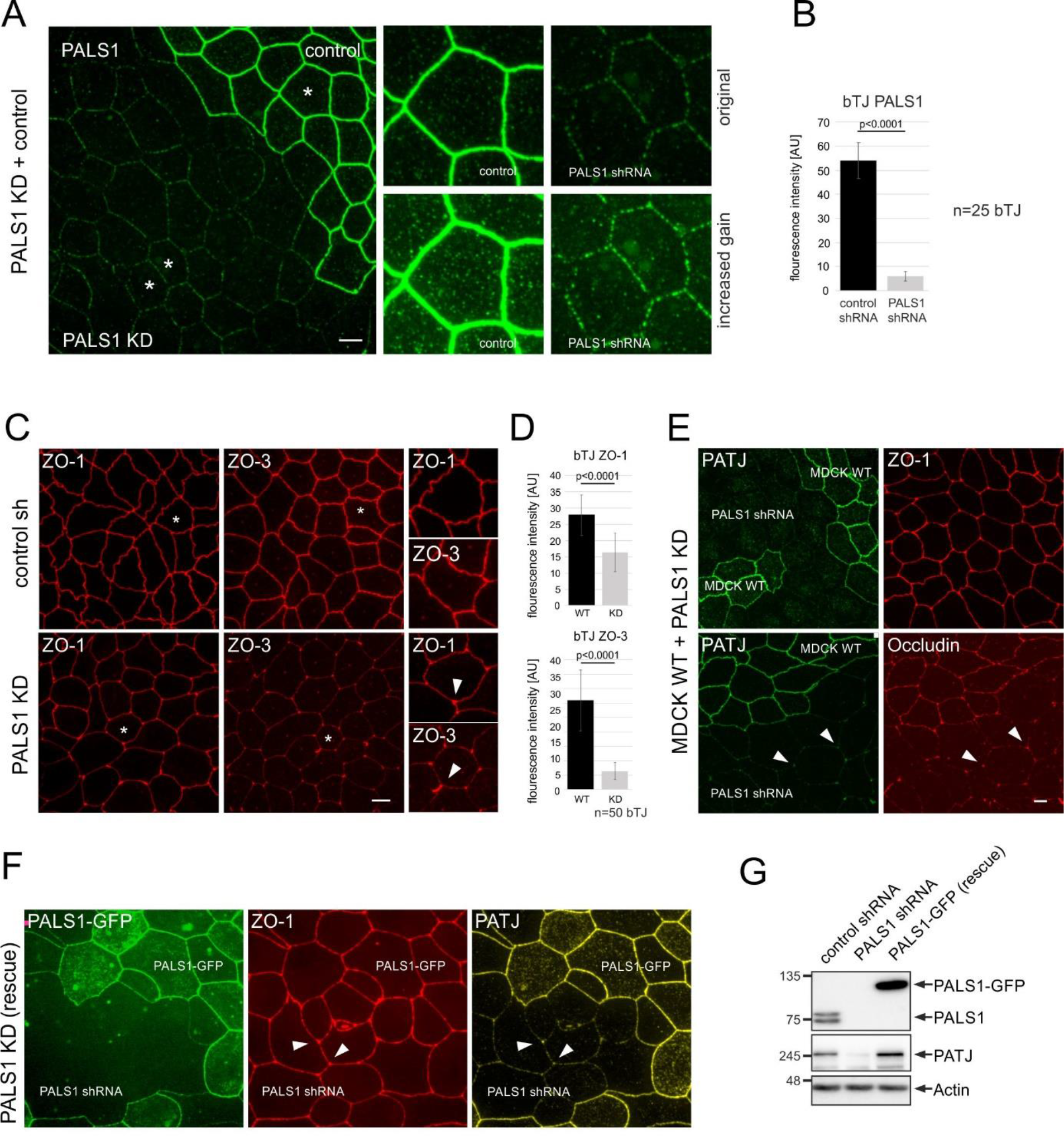
PALS1 regulates the distribution of TJ proteins and PATJ in a dose dependent maner. (A) IF analysis of co-cultures of PALS1 shRNA knockdown cells (PALS1 KD) and control cells stably expressing a non- targeting shRNA (control) show a strong reduction in PALS1 expression. Residual PALS1 produces a fine punctate pattern along the bicellular junctions (details: original and increased gain). Scale bar: 5 µm. (B) Quantitative evaluation of the PALS1 KD efficiency. (C) IF analyses of ZO proteins in PALS1 KD and WT cells. PALS1 downregulation leads to a strong reduction of ZO-1 and ZO-3 in bicellular TJ, leading to a “dot-like” pattern, like PALS1 KO cells. Scale bar: 5 µm. (D) Quantification of the IF data (graph on the right) confirms the strong reduction of PALS1 in bicellular TJ (bTJ). (E) IF analyses of co-cultures of MDCK WT and PALS1 KD cells. *Upper panel:* Staining of ZO-1 (red) and PATJ (green) revealed a strong reduction of PATJ in PALS1 KD cell patches. *Lower panel:* In MDCK WT cells PATJ (green) co-localizes with Occludin (red). In PALS1 KD patches, PATJ and Occludin co-localize in a “dot- like” pattern, due to the disrupted distribution of both proteins along bicellular TJ. Scale bar: 5 µm. (F) Rescue experiment. The disapearence of PATJ from bicellular junctions in PALS1 KD cells is completely rescued by the re- expression of PALS1-GFP (green). Arrowheads mark “dot-like” structures of ZO-1 (red) and PATJ (yelleow) in PALS1 KD cells. (G) Western blot analysis of the control shRNA line, the PALS1 shRNA line, and the PALS1-GFP rescue line (PALS1-GFP expressed in PALS1 KD line). Note the strong reduction of PATJ in the PALS1 KD line, which is rescued by re-expression of PALS1-GFP.

## Discussion

In this study, we analyzed the functions of PALS1 in MDCKII cells. Genetic inactivation of the *PALS1* gene using CRISPR/Cas9 resulted in an increased formation of multiple- and no-lumen phenotypes in 3D cyst assays and a strong increase in paracellular permeability in cysts derived from PALS1 KO cells. PALS1 KO cells that formed single-lumen cysts also showed an increase in paracellular permeability, whereas single- and multiple-lumen cysts of MDCK WT cells maintained a tight paracellular diffusion barrier for solutes – at least up to 10 kDa. This is in line with earlier studies [6] und suggests that PALS1 is involved in the regulation of the paracellular transport of ions and solutes in epithelial cells and that the formation and function of TJs is not directly coupled to the establishment of a single apical luminal domain.

Strikingly, PALS1-deficient cells revealed a significant reduction in TJ proteins (ZO-1, ZO-3 and Occludin) along bicellular junctions, accompanied by an accumulation of these proteins at tricellular junctions, resulting in a “dot-like” (or “spike-like”) distribution pattern. Remarkably, this discontinuous distribution was preserved in bi- and tricellular junctions that were formed between WT and PALS1 KO cells. In other words: PALS1 KO cells that formed cell-cell contacts with cells that express PALS1 (MDCK WT cells) maintained a disrupted TJ distribution, indicating that PALS1 on one side of cell-cell contacts is unable to trigger full TJ formation in adjoining cells. A similar observation was made in co-cultures of WT and PALS1 KD cells. Thus, together the data suggests, first, that PALS1 controls the proper lateral distribution of TJ proteins along the bicellular contacts in a dose-dependent manner, and second, that the continuous lateral ZO-1 distribution of intercellular TJs is directly linked to the cell-autonomous expression of PALS1.

Interestingly, in PALS1 KO and KD cells PATJ was lost from bicellular TJs and instead co-localized with TJ proteins such as ZO-1 and Occludin at tricellular junctions. This indicates that PATJ localization is dependent on PALS1 and suggests that PALS1 acts upstream of PATJ. However, the situation is probably more complicated. Earlier studies identified the L27N domain of PALS1 as the PATJ binding module that targets PALS1 to TJs [21], suggesting that membrane targeting of PALS1 requires the presence of PATJ. Moreover, we show that PALS1 deletion mutants lacking the L27N domain do not rescue the ZO-1 distribution phenotype of PALS1 KO cell lines, indicating a crucial role for PATJ, or at least of the PALS1 PATJ-binding module, in regulating the distribution of ZO-1 (and other TJ proteins). These data are in agreement with earlier studies in Caco2 cells showing that PATJ regulates the localization of Occludin and ZO-3 to TJs [22]. Moreover, it has recently been demonstrated that MDCK cells that express a PATJ variant lacking the PALS1- binding module (ΔL27-PATJ) failed to form a continuous TJ belt due to a reduced spreading dynamics of ZO-1-condensates[23]. Intriguingly, the disrupted ZO-1 distribution along bicellular junctions in ΔL27-PATJ MDCK cells strongly resembles the “dot-like” pattern that we observed in MDCK PALS1 KO und KD cell lines[23]. Together these observations indicate that PALS1 and PATJ are dependent on one another and that a close functional coupling between PALS1 and PATJ is required for the formation of a continuous and functional TJ belt in epithelial cells. The strong interconnection between PALS1 and PATJ is also supported by recent super-resolution imaging studies that demonstrate a high degree of co-localization of these proteins at the apical-lateral border [4]. Thus, our observations, those by *Pombo-Garcia et al*, and earlier studies by the groups of *Margolis* and *Le Bevic* indicate that the lateral continuous belt formation of ZO-1 (and other TJ proteins) may be predominantly organized *via* a PALS1-PATJ axis[6, 21–23].

Proteins of the ZO-family are necessary for the formation of TJ strands [24, 25]. The loss of ZO-1 and ZO-2 in 3D MDCKII cysts is associated with a dramatic increase in paracellular permeability to water, solutes, and ions[24]. Thus, the ZO-1/ZO-2 double KO cell lines show a phenotype quite similar to MDCK cells lacking PALS1 (this study) and cells lacking the PALS1 binding site in PATJ (ΔL27-PATJ[23]). These findings underline the essential role of ZO proteins in regulating the paracellular transport in epithelia [26, 27] and suggest that PALS1 – most likely in concert with PATJ – acts upstream of ZO-1 and other components the TJs. PALS1 also interacts with partitioning-defective 6 (PAR6) [18, 19], which is part of the PAR complex that controls apicobasal cell polarization in concert with the CRB complex [19]. However, we observed that PALS1 without the PAR6 binding site (ΔECR-PALS1) can re-establish the lateral ZO-1 distribution defect of PALS1 KO cells, indicating that the PALS1-PAR6 interaction is less relevant for the continuous distribution of a range of TJ proteins including ZO-1, ZO-3 and Occludin.

Both, PALS1 and PATJ, interact not only with polarity and TJ proteins. Recent proximity labeling studies have identified PALS1 as a core component of a very complex network in the vertebrate apical marginal zone that includes both known (*e.g.* PATJ and LIN7) and novel binding partners of PALS1 (*e.g.* HOMER proteins) that may provide links to TJ proteins or regulate epithelial functions beyond the formation of cell-cell contacts[5]. Strikingly, and possibly of particular importance, PALS1 but also PATJ interact with components and regulators of the Hippo pathway. PATJ, for example, binds not only to the TJ proteins ZO-3 and Claudin1 but also to the AMOT protein family and KIBRA, both of which are important regulators of Hippo-YAP/TAZ signaling [28–31]. In addition, both PALS1 and PATJ were reported to bind to YAP and TAZ, the transcriptional coactivators of Hippo signaling [14, 28, 32]. Interestingly, the loss of TAZ in the kidney results in a similar phenotype[33–36] as the loss of PALS1, LIN7, or CRB3a[8–11], providing further evidence that components of the CRB complex are closely linked to Hippo- signaling[37–39].

The epithelia of the nephron segments in the kidney ensure the excretion of noxious substances, the concentration of the primary ultra-filtrate into the final urine as well as the recycling and resorption of numerous nutrients and organic and inorganic ions. All these functions require a fine-tuned control of the paracellular transport processes, which in turn depends on the nephron segment-specific expression of TJ components, for example the numerous members of the Claudin family [27, 40, 41]. In other words: minor changes in TJ formation dynamics, maintenance or in the distribution of central TJ proteins may have very severe consequences *in vivo*. The dysfunctional lateral distribution of TJ proteins in PALS1 KO and KD cells and the strong phenotype of Pals1 haplo-insufficient mice in which already a partial Pals1 depletion caused severe defects – although the well-defined apicobasal distribution of important proteins is in principle preserved – supports this assumption[11, 12]. Further research will be necessary to clarify to what extent even minor modifications of the “tightness of TJs” contribute to renal pathomechanisms.

## Author contributions

AG performed most of the experimental work, supported by SK, MB, VH and AMK for sub-aspects. PN, MPK, AL and KE provided cell lines, antibodies, and expertise. SG and AL performed and analyzed all PALS1 shRNA experiments and UH performed the ultrastructural analyses. AG, HP and TW account for the experimental design. AG and TW prepared the manuscript.

## Acknowledgements

We would like to thank Dr. Alf Honigmann (MPI Dresden) for providing us details of the permeability protocol. We would like to thank Karin Gäher for excellent technical support. The work was supported by DFG grant (WE 2550/4- 1 of SPP1782) to TW and by the Graduate School of the Cells-in-Motion Cluster of Excellence (EXC 1003 – CiM to AG). This research was supported by the Ministry of Education, Singapore, under its Academic Research Fund Tier2 (MOE-T2EP30121-0019) to AL.

## Competing Interests

The authors have no relevant financial or non-financial interests to disclose.

## Data availability

The data and tools that support the findings of this study are available from the corresponding author upon reasonable request.

## Supplemental Data

### Table of content

Suppl. Table ST1: Primers and oligonucleotides.

Suppl. Table ST2: Primary antibodies.

Suppl. Table ST3: Secondary antibodies.

Suppl. Fig. S1: Generation of and validation of MDCK PALS1 knockout cell lines.

Suppl. Fig. S2: Permeability assays of MDCK wildtype and PALS1 KO.

Suppl. Fig. S3: Monolayers lacking PALS1 show an altered distribution of tight junction proteins.

Suppl. Fig. S4: The continuous lateral TJ distribution is linked to cellular PALS1 expression.

Suppl. Fig. S5: PALS1 levels control the lateral distribution of TJ proteins ZO-1 and ZO-3.

Suppl. Fig. S6: Rescue by PALS1-GFP re-establishes PATJ expression and localization.

**Suppl. Figure S1:**
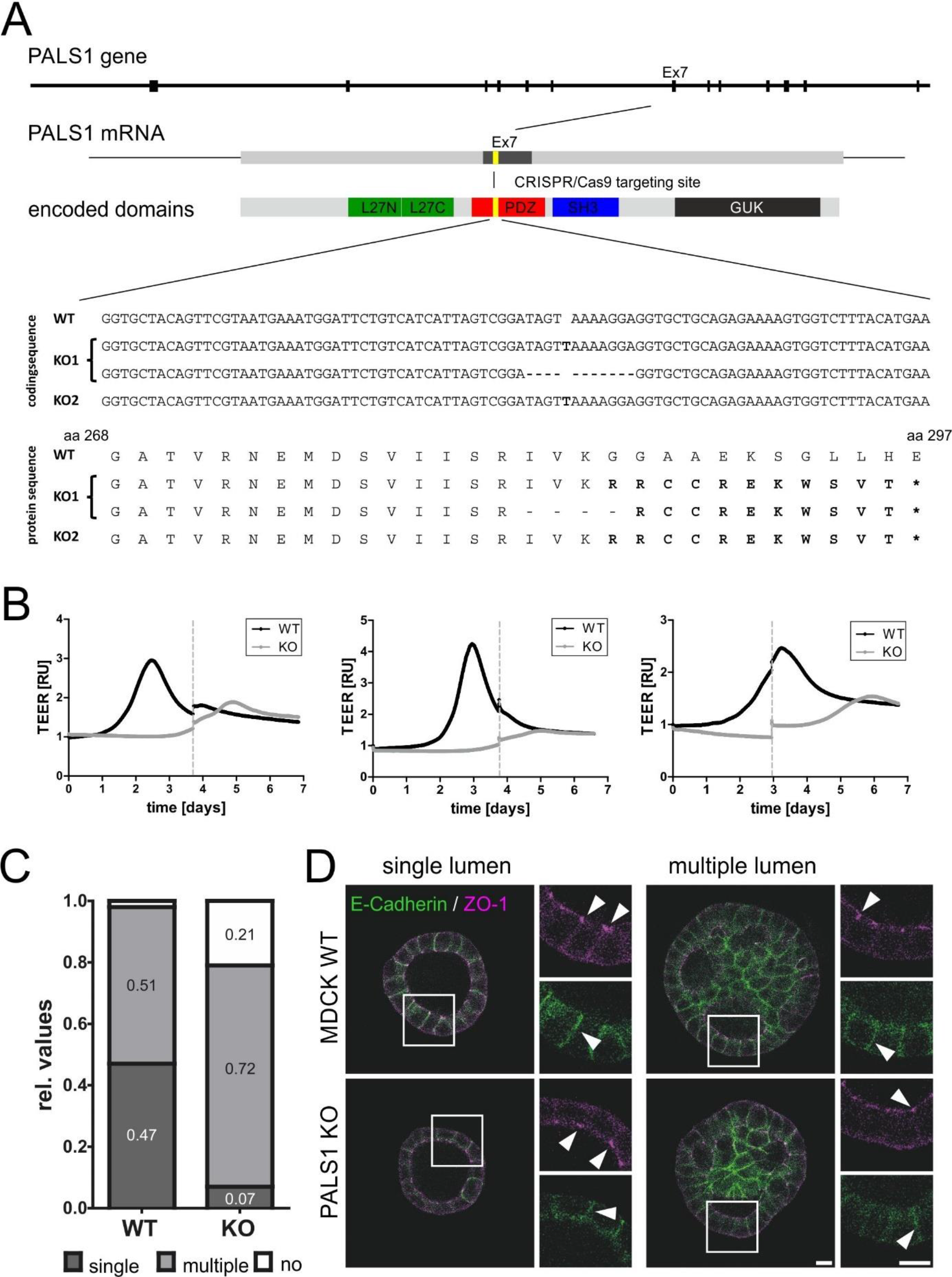
Generation of and validation of MDCK PALS1 knockout cell lines. (A) Scheme: The canine *PALS1* gene was inactivated by CRISPR/Cas9 using a guidance sequence (gDNA that tragets coding exon 7, which encodes parts of the PALS1 PDZ domain with. Sequencing revealed a 11 bp deletion and 1 bp insertion for knockout clone 1 (KO1) and an 1 bp insertion in both allels for knockout clone 2 (KO2). Both CRISPR/Cas9 mediated changes resulte in an premature trunction of the PALs1 encoding amino acid sequence due to alternative stop (*) codons. (B) Graphs of three independent TEER measurements of MDCK WT (black) WT and PALS1 KO cell lines (grey). Directly after seeding, measurements were taken for the next seven days at an interval of every 5 min. WT cells show resistancy maximum peak between (between day 2.5. and 3.5). The maximum resistance peak in in KO cells is strongly decreased. The resistance maximum peaks of KO cells have a delay of approximately 2 days. The dashed line indicate the interruption of the measurement due to change of medium, RU: relative units. (C) Lumen formation of MDCK PALS1 KO and WT cells. PALS1 KO cells show an increased number of multiple-, and no-lumen cyst phenotypes. Y-axis is given in relative values (1 ≙ 100% of the counted cysts). For the experiment we analyzed MDCKII WT (n=242) and PALS1 KO (n= 177) cyst of three independent experiments (N=3). Chi square test showed a significant difference in the distribution of lumen phenotype between WT and KO cysts (*χ*^2^=48.91; df=2; p ≤ 0.001) (D) IF analyses of 3D cysts. Staining of ZO-1 and E-Cadherin in single- and multiple lumen 3D cysts demonstrate similar distribution of these proteins in PALS1 KO and WT cysts. Scale bar 10 µm.

**Suppl. Figure S2:**
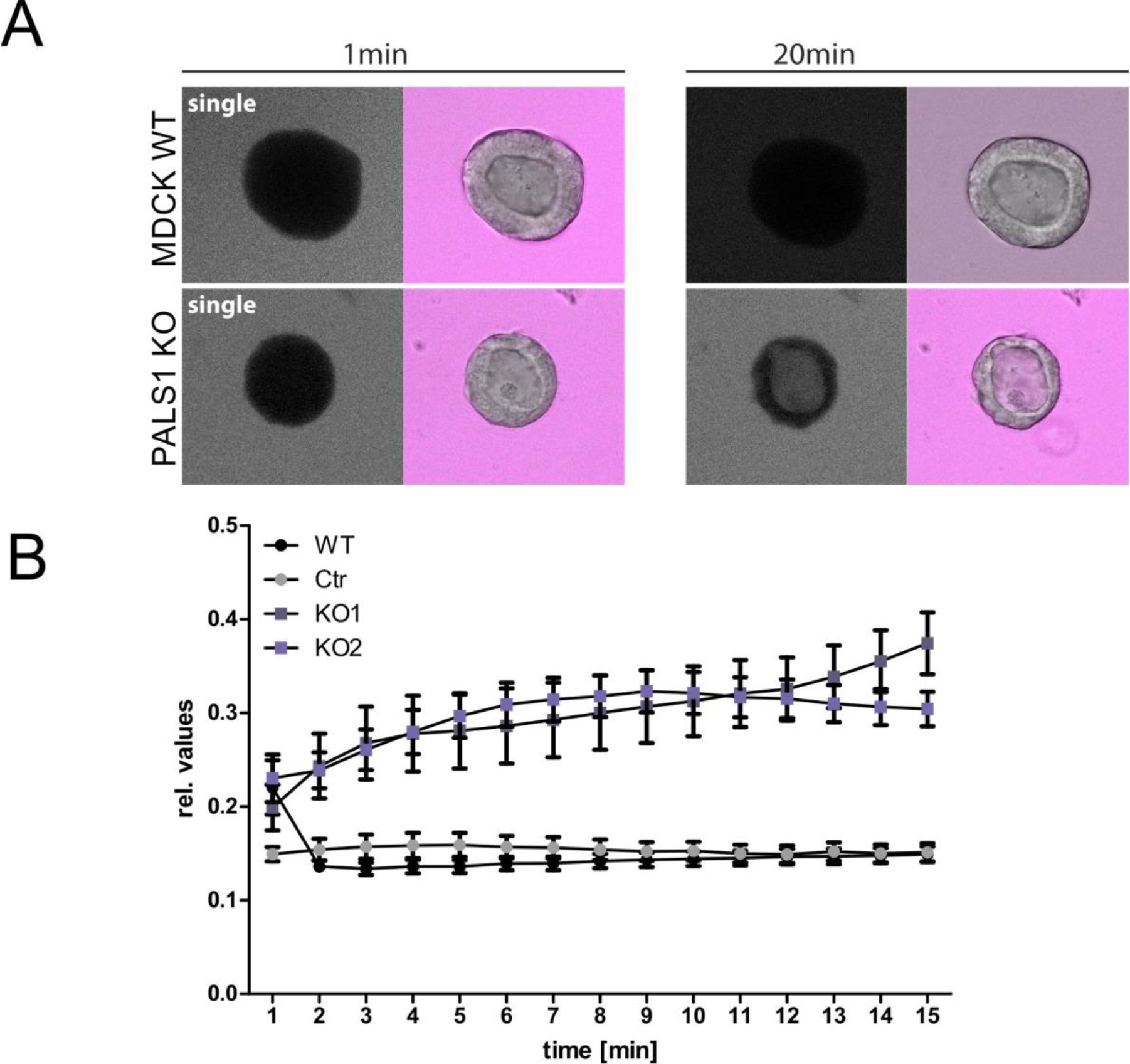
Permeability assays of MDCK wildtype and PALS1 KO. (A) Permeability assays. Images show an accumulation of Alexa Fluor™ 647-labelled 10kDa dextran only in in the lumen of PALS1 KO cysts after 20 min (N=3). (B) Time course of the permeability assay. Cyst permeability assays were performed for 20 min. Signal intensity in the lumen was normalized to the surrounding signal in the medium. No uptake was detected in WT cysts. In contrast, PALS1 KO cysts show a strong increase of labelled dextran inside the cyst lumen over time.

**Suppl. Fig. S3:**
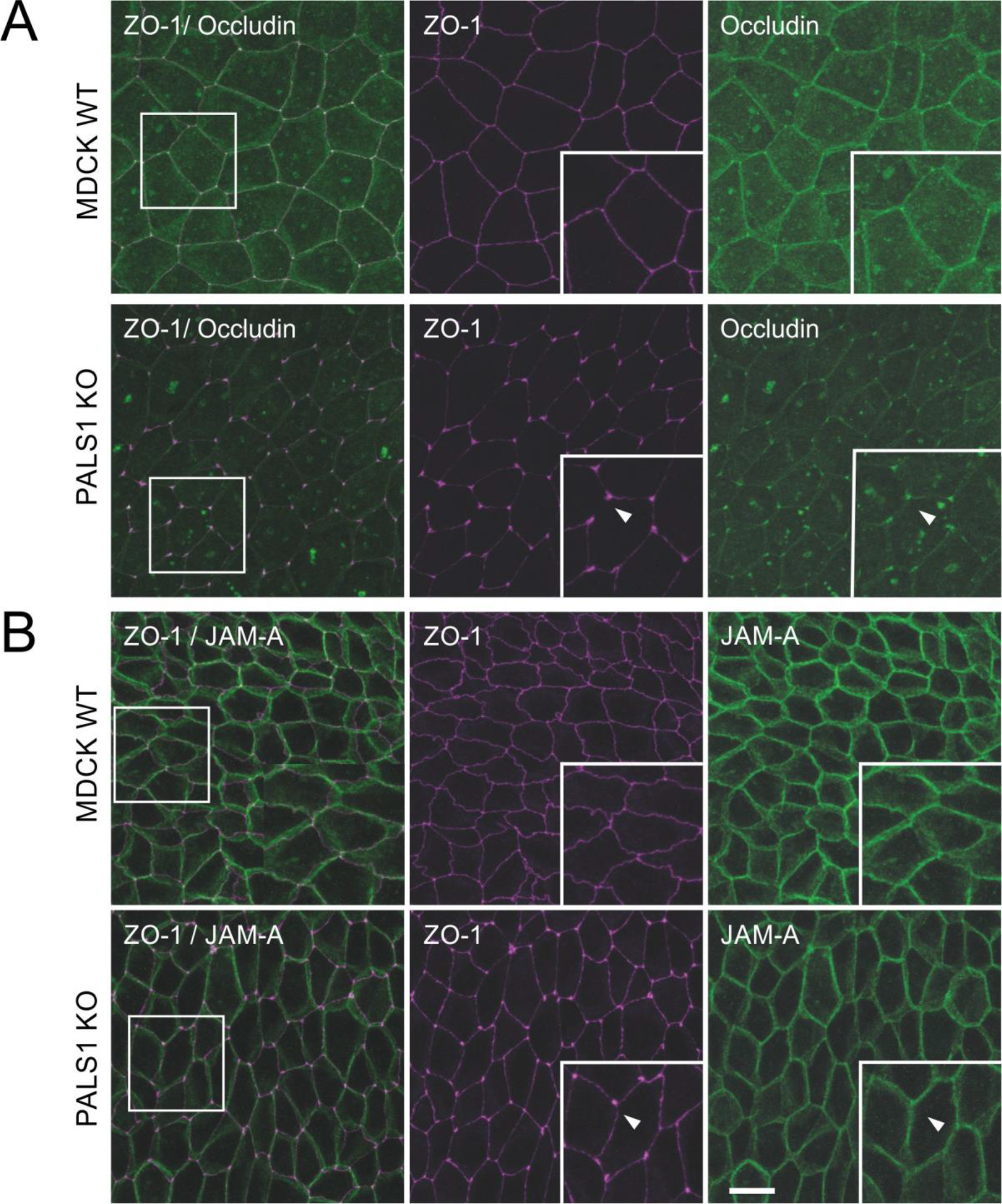
Monolayers lacking PALS1 show an altered distribution of tight junction proteins. (A) IF analyses of MDCK WT and PALS1 KO cells. MDCKII cells were co-stained with antibodies against ZO-1 (magenta) and Occludin (green). In WT cells, ZO-1 and Occludin showed a “chicken-wire-like” distribution. In PALS1 KO cells TJ proteins Occludin and ZO-1 are strongly reduced along bicellular junction leading to a “dot-like” pattern, accompanied by a tricellular accumulation of these proteins (arrow heads). (B) JAM-A showed a similar “chicken-wire-like” distribution in both WT and PALS1 KO monolayers. Images show a maximum intensity projection. Scale bar: 10 µm

**Suppl. Fig. S4:**
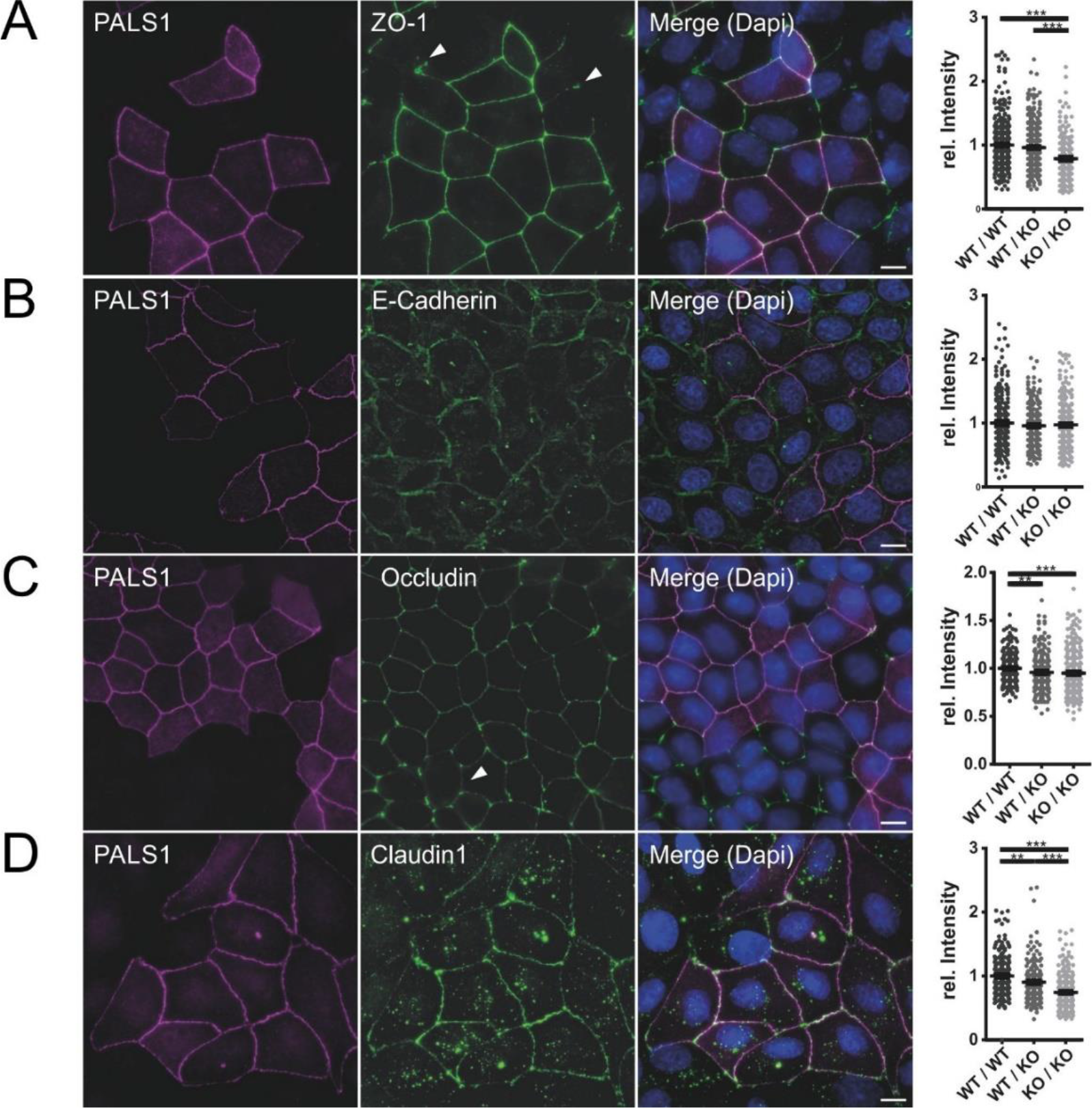
The continuous lateral TJ distribution is linked to cellular PALS1 expression. Co-cultures of MDCKII WT and Pals1 KO cells grown on glass cover slips were stained against PALS1, ZO-1, E-Cadherin, Occludin, or Claudin1 to analyze shared WT-WT, WT-KO, and KO-KO cell contacts. Images show a maximum intensity projection. Brightness and contrast were increased equally for all images. Cells were mixed in a ratio of 1:1. (A) IF study for PALS (magenta) and ZO-1 (green): Staining revealed a strong reduction of ZO-1 in PALS1 KO cell patches, with reduced signals along the bicellularand accmulations in tricellularcontacts (arrow heads). Signals between WT-WT (n=471), WT-KO (n=491) and KO-KO (n=206) contacts were quantified and normalized to the mean intensity of the bicellular WT-WT junctions (graph, right of the panel). (B) IF study for PALS (magenta) and E-Cadherin (green): E-Cadherin shows a normal junctional distribution in both MDCK WT and PALS1 KO cells patches. No significant signal intensity differences between WT and KO cells were detectable. WT-WT (n=306), WT-KO (n=281) and KO-KO (n=288) (C) IF study for PALS (magenta) and Occludin (green): Occludin showed a similar pattern as ZO-1 (see panel A), leading to reduced levels along bi- and accumulations within tricellular contacts of PALS1 KO patches. WT-WT (n=256), WT-KO (n=254) and KO-KO (n=288) (D) IF study for PALS (magenta) and Claudin1: The Claudin1 showed a continuous distribution and co-localization with PALS1. This distribution is completely disrupted in PALS1 KO cells. WT-WT (n=223), WT-KO (n=223) and KO-KO (n=263). Scale bar: 10 µm.***≙ p ≤ 0.001; **≙ p ≤ 0.01

**Suppl. Fig. S5:**
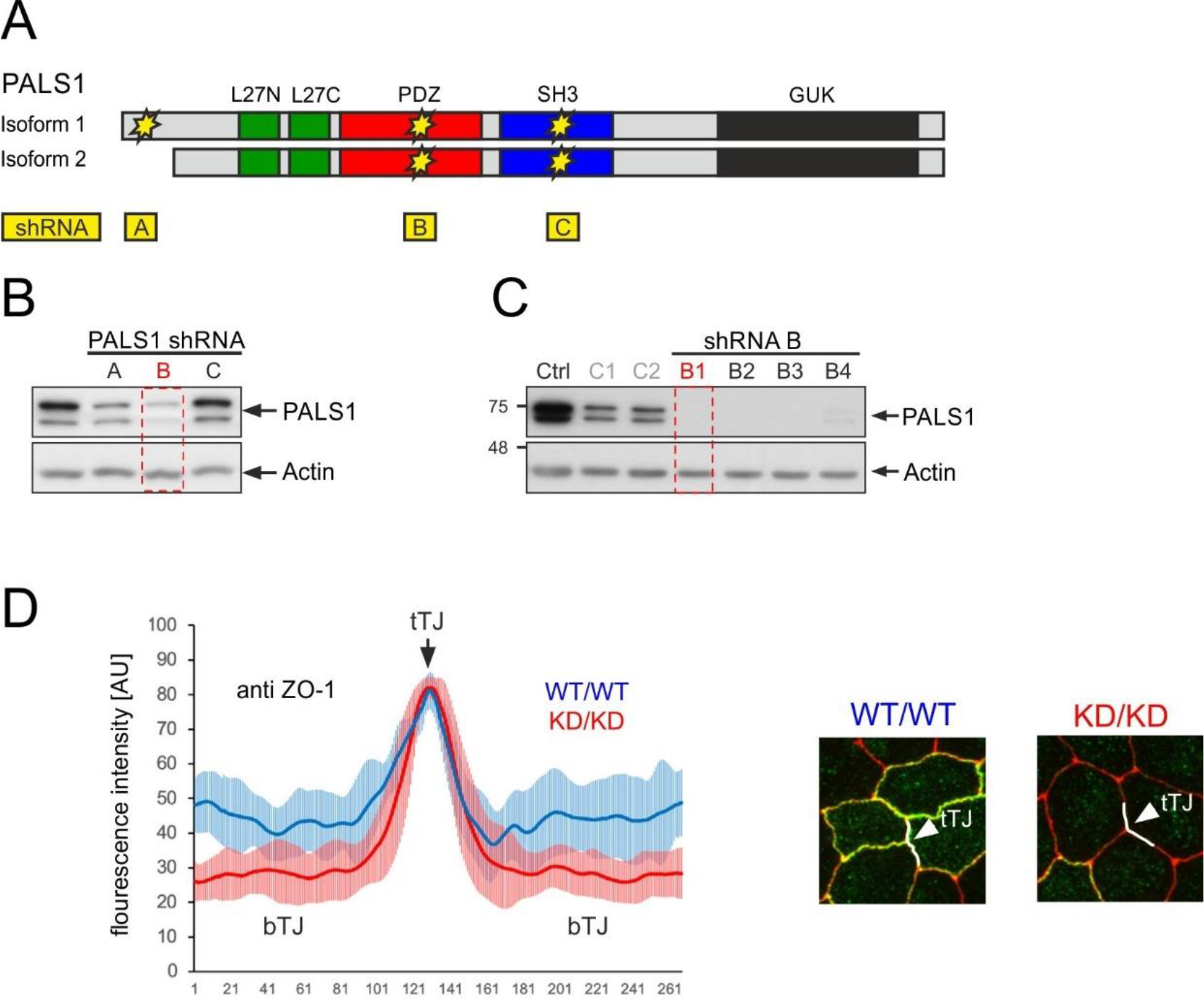
PALS1 levels control the lateral distribution of TJ proteins ZO-1 and ZO-3. (A) Scheme: Domain architecture of PALS1 isoforms (green: L27 domain, red: PDZ domain, blue SH3 domain, Black: GUK domain) with shRNA targeting sites for PALS1 silencing. Asterisks mark the positions of shRNAs (A, B, C) designed to target the PALS1 mRNA. (B) Western blots of cell lysates of MDCK cell pools stably transfected with shRNAs A, B, or C. The shRNA B was the most efficient shRNA. (C) Western blots of clonal cell lines expressing shRNA B. Clone B1 was used for all further analyses (D) IF analysis PALS1 KD and control cells stably expressing a non-targeting shRNA. Quantification of the IF data confirms the strong reduction of PALS1 in bicellular TJ (bTJ).

**Suppl. Fig. S6:**
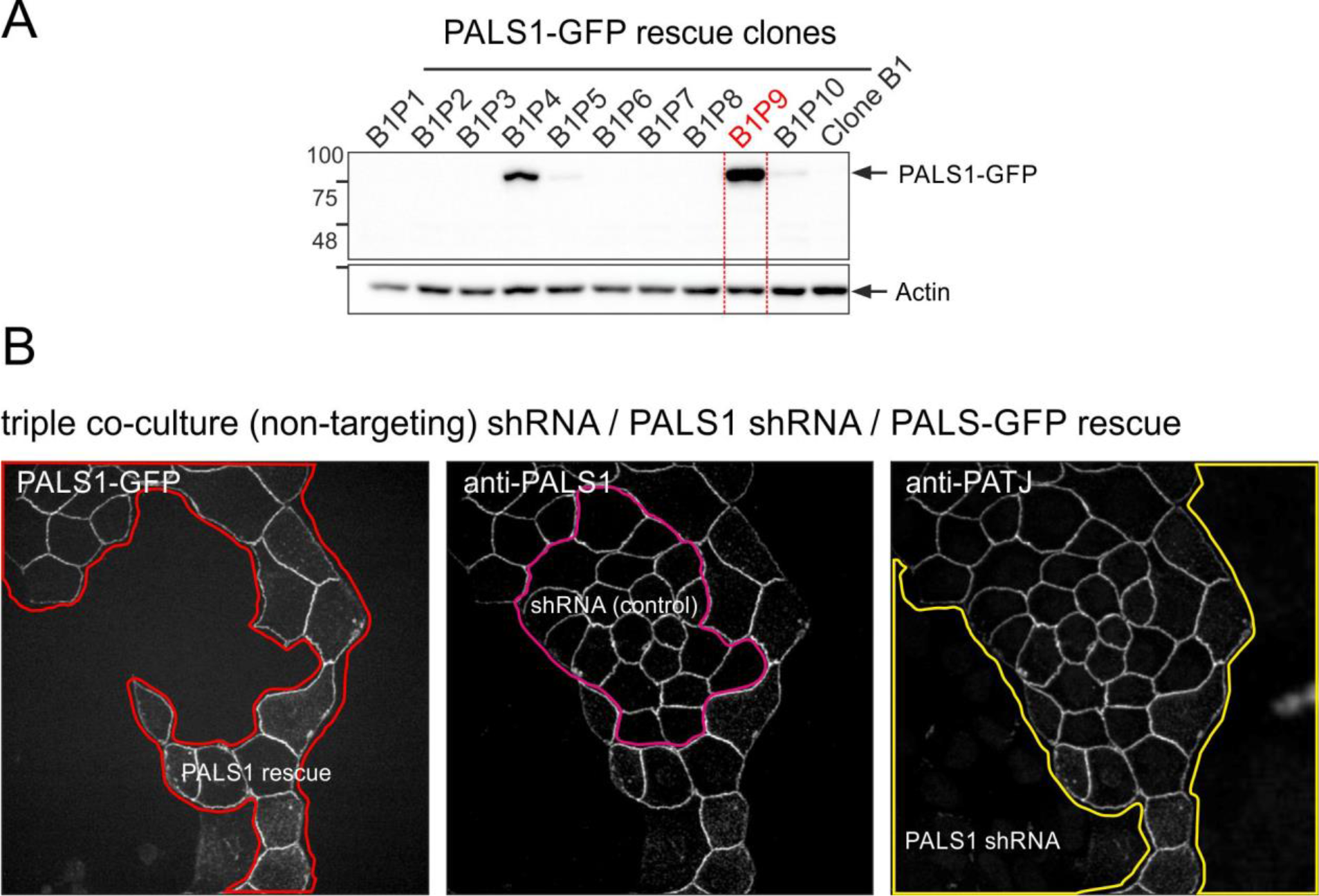
Rescue by PALS1-GFP re-establishes PATJ expression and localization. (A) MDCK cell clone B1P9 showed a strong expression of PALS1-GFP in a PALS1 KD cells (clone B1). (B) IF analyses of triple co-cultures. Mixed monolayers of PALS1 KD cells (clone B1), control cells (stably expressing a non-targeting shRNA), and rescue cell line (stable expressing PALS1-GFP, show that PATJ expression and localization depends on PALS1, as only PALS1-GFP expressing cells (red patch), and shRNA control cells (magenta patch) show PATJ positive cells.

**Suppl. Table ST1:**
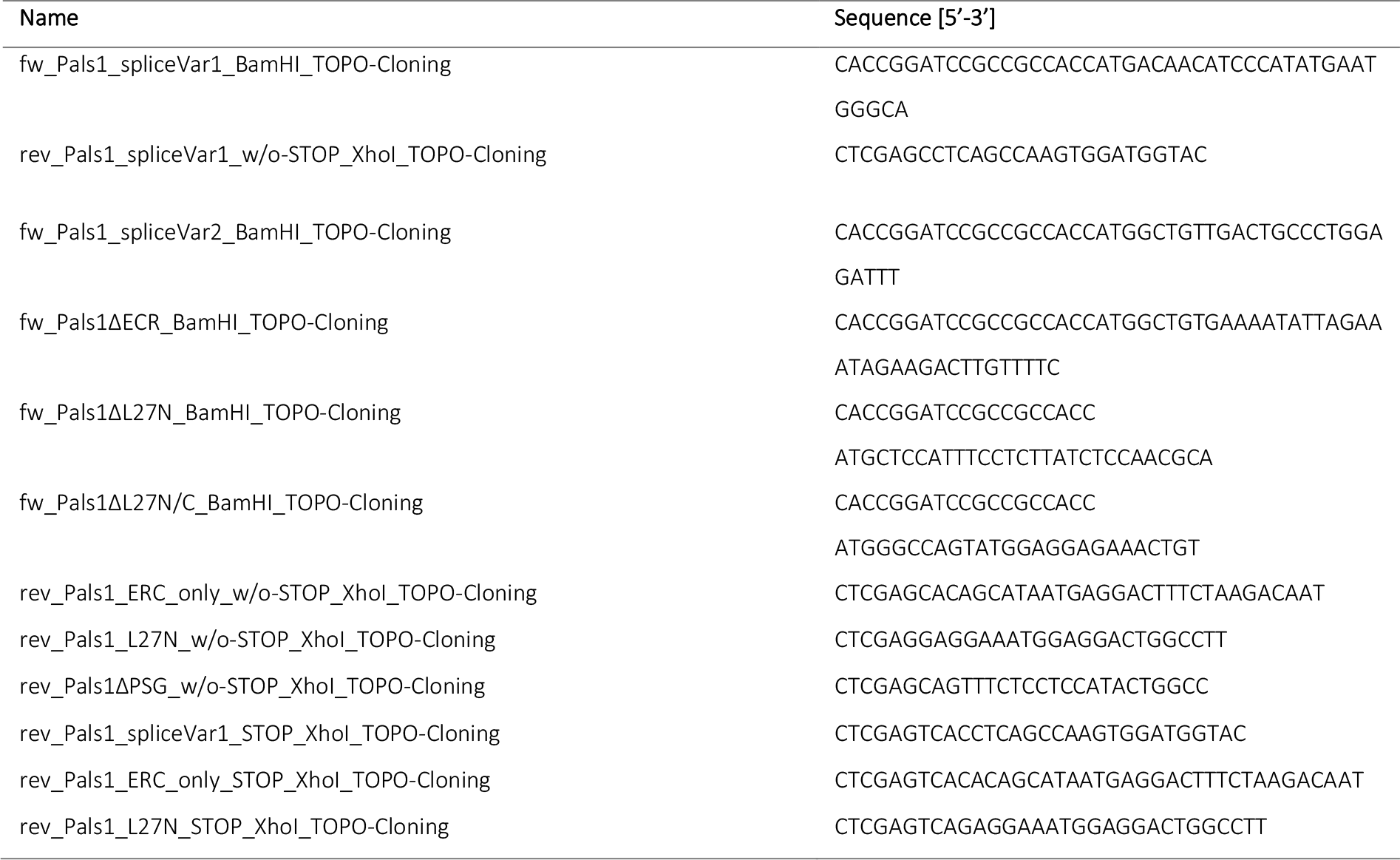
Primers and oligonucleotides.

**Suppl. Table ST2:**
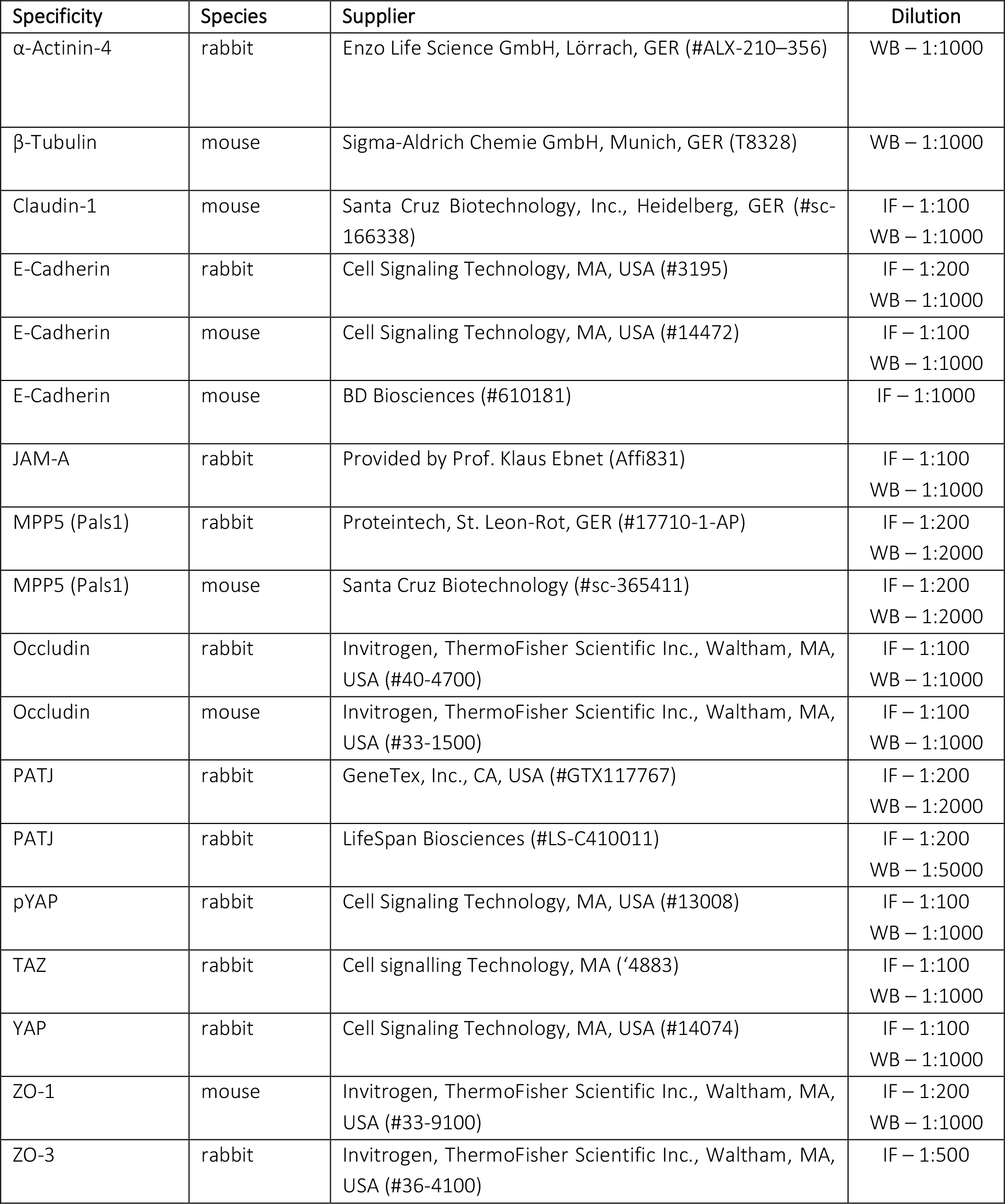
Primary antibodies.

| Specificity | Species | Supplier | Dilution |
| --- | --- | --- | --- |
| α-Actinin-4 | rabbit | Enzo Life Science GmbH, Lörrach, GER (#ALX-210-356) | WB – 1:1000 |
| β-Tubulin | mouse | Sigma-Aldrich Chemie GmbH, Munich, GER (T8328) | WB – 1:1000 |
| Claudin-1 | mouse | Santa Cruz Biotechnology, Inc., Heidelberg, GER (#sc-166338) | IF – 1:100<br>WB – 1:1000 |
| E-Cadherin | rabbit | Cell Signaling Technology, MA, USA (#3195) | IF – 1:200<br>WB – 1:1000 |
| E-Cadherin | mouse | Cell Signaling Technology, MA, USA (#14472) | IF – 1:100<br>WB – 1:1000 |
| E-Cadherin | mouse | BD Biosciences (#610181) | IF – 1:1000 |
| JAM-A | rabbit | Provided by Prof. Klaus Ebnet (Affi831) | IF – 1:100<br>WB – 1:1000 |
| MPP5 (Pals1) | rabbit | Proteintech, St. Leon-Rot, GER (#17710-1-AP) | IF – 1:200<br>WB – 1:2000 |
| MPP5 (Pals1) | mouse | Santa Cruz Biotechnology (#sc-365411) | IF – 1:200<br>WB – 1:2000 |
| Occludin | rabbit | Invitrogen, ThermoFisher Scientific Inc., Waltham, MA, USA (#40-4700) | IF – 1:100<br>WB – 1:1000 |
| Occludin | mouse | Invitrogen, ThermoFisher Scientific Inc., Waltham, MA, USA (#33-1500) | IF – 1:100<br>WB – 1:1000 |
| PATJ | rabbit | GeneTex, Inc., CA, USA (#GTX117767) | IF – 1:200<br>WB – 1:2000 |
| PATJ | rabbit | LifeSpan Biosciences (#LS-C410011) | IF – 1:200<br>WB – 1:5000 |
| pYAP | rabbit | Cell Signaling Technology, MA, USA (#13008) | IF – 1:100<br>WB – 1:1000 |
| TAZ | rabbit | Cell signalling Technology, MA ('4883) | IF – 1:100<br>WB – 1:1000 |
| YAP | rabbit | Cell Signaling Technology, MA, USA (#14074) | IF – 1:100<br>WB – 1:1000 |
| ZO-1 | mouse | Invitrogen, ThermoFisher Scientific Inc., Waltham, MA, USA (#33-9100) | IF – 1:200<br>WB – 1:1000 |
| ZO-3 | rabbit | Invitrogen, ThermoFisher Scientific Inc., Waltham, MA, USA (#36-4100) | IF – 1:500 |

**Suppl. Table ST3:** Secondary antibodies.

| Specificity | Species | Supplier | Dilution |
| --- | --- | --- | --- |
| Alexa Fluor® 488-α-mouse IgG | goat | Invitrogen Life Technologies, Darmstadt, GER | IF 1:1000 |
| Alexa Fluor® 594-α-mouse IgG | goat | Invitrogen Life Technologies, Darmstadt, GER | IF 1:1000 |
| Alexa Fluor® 488-α-rabbit IgG | goat | Invitrogen Life Technologies, Darmstadt, GER | IF 1:1000 |
| Alexa Fluor® 594-α-rabbit IgG | goat | Invitrogen Life Technologies, Darmstadt, GER | IF 1:1000 |
| Alexa Fluor® 647-α-rabbit IgG | goat | Invitrogen Life Technologies, Darmstadt, GER | IF 1:1000 |
| Janelia Fluor® 646 HALO tag ligand | - | Promega GmbH, Walldorf, GER | IF 1:500 |
| Peroxidase AffiniPure Goat Anti-mouse IgG | goat | Jackson ImmunoResearch Europe Ltd., Ely, UK | WB 1:1000 |
| Peroxidase AffiniPure Goat Anti-rabbit IgG | goat | Jackson ImmunoResearch Europe Ltd., Ely, UK | WB 1:1000 |

## Notes

### Competing Interest Statement

The authors have declared no competing interest.

